# Sex determination gene *transformer* regulates the male-female difference in *Drosophila* fat storage via the adipokinetic hormone pathway

**DOI:** 10.1101/2021.07.20.453098

**Authors:** Lianna W. Wat, Zahid S. Chowdhury, Jason W. Millington, Puja Biswas, Elizabeth J. Rideout

**Affiliations:** Department of Cellular and Physiological Sciences, Life Sciences Institute, The University of British Columbia

**Keywords:** *Drosophila*, fat storage, triglyceride, adipokinetic hormone, sex determination, *transformer*, lifespan, fertility

## Abstract

Sex differences in whole-body fat storage exist in many species. For example, *Drosophila* females store more fat than males. Yet, the mechanisms underlying this sex difference in fat storage remain incompletely understood. Here, we identify a key role for sex determination gene *transformer* (*tra*) in regulating the male-female difference in fat storage. Normally, a functional Tra protein is present only in females, where it promotes female sexual development. We show that loss of Tra in females reduced whole-body fat storage, whereas gain of Tra in males augmented fat storage. Tra’s role in promoting fat storage was largely due to its function in neurons, specifically the Adipokinetic hormone (Akh)-producing cells (APCs). Our analysis of Akh pathway regulation revealed a male bias in APC activity and Akh pathway function, where this sex-biased regulation influenced the sex difference in fat storage by limiting triglyceride accumulation in males. Importantly, Tra loss in females increased Akh pathway activity, and genetically manipulating the Akh pathway rescued Tra-dependent effects on fat storage. This identifies sex-specific regulation of Akh as one mechanism underlying the male-female difference in whole-body triglyceride levels, and provides important insight into the conserved mechanisms underlying sexual dimorphism in whole-body fat storage.

## INTRODUCTION

In animals, stored fat provides a rich source of energy to sustain basal metabolic processes to survive periods of nutrient scarcity, and to support reproduction (Heier and Kühnlein, 2018; Heier et al., 2021; Walther and Farese, 2012). The main form of stored fat is triglyceride, which is deposited within specialized organelles called lipid droplets (Kühnlein, 2012; Murphy, 2001; Thiele and Spandl, 2008). Lipid droplets are found in many cell types throughout the body, but the main organ responsible for triglyceride storage is the adipose tissue (Murphy, 2001). The amount of triglyceride in the adipose tissue is regulated by many factors; however, one important factor that influences an individual’s whole-body fat level is whether the animal is female or male (Karastergiou et al., 2012; Power and Schulkin, 2008; Sieber and Spradling, 2015; Wat et al., 2020). Typically, females store more fat than males. In mammals, females store approximately 10% more body fat than males (Jackson et al., 2002; Karastergiou et al., 2012; Womersley and Durnin, 1977). Female insects, on the other hand, can store up to four times more fat than males of the same species (Lease and Wolf, 2011) and break down fat more slowly than males when nutrients are scarce (Wat et al., 2020). These male-female differences in fat metabolism play a key role in supporting successful reproduction in each sex: females with reduced fat storage often show lower fecundity (Buszczak et al., 2002; Sieber and Spradling, 2015) whereas males with excess fat storage generally show decreased fertility (Grönke et al., 2005; Wat et al., 2020). Given that fat storage also influences diverse phenotypes such as immunity and lifespan (DiAngelo and Birnbaum, 2009; Gáliková and Klepsatel, 2018; Johnson and Stolzing, 2019; Kamareddine et al., 2018; Liao et al., 2021; Roth et al., 2018; Suzawa et al., 2019), the sex-specific regulation of fat storage has implications for several life history traits. Yet, the genetic and physiological mechanisms that link biological sex with fat storage remain incompletely understood in many animals.

Clues into potential mechanisms underlying the sex difference in fat storage have emerged from studies on the regulation of triglyceride metabolism in *Drosophila*. While many pathways impact whole-body triglyceride levels (Ballard et al., 2010; Bjedov et al., 2010; Broughton et al., 2005; Choi et al., 2015; DiAngelo and Birnbaum, 2009; Francis et al., 2010; Ghosh and O’Connor, 2014; Grönke et al., 2010; Heier and Kühnlein, 2018; Heier et al., 2021; Hentze et al., 2015; Kamareddine et al., 2018; Kang et al., 2017; Kubrak et al., 2020; Lee et al., 2019; Lehmann, 2018; Luong et al., 2006; Rajan and Perrimon, 2012; Roth et al., 2018; Scopelliti et al., 2019; Sieber and Spradling, 2015; Song et al., 2014, 2017; Suzawa et al., 2019; Teleman et al., 2005; Texada et al., 2019), the Adipokinetic hormone (Akh; FBgn0004552) pathway plays a central role in regulating whole-body fat storage and breakdown (Heier and Kühnlein, 2018; Heier et al., 2021; Lehmann, 2018). Akh is synthesized as a preprohormone in the Akh-producing cells (APCs), and is subsequently cleaved by proprotein convertases to produce active Akh (Lee and Park, 2004; Noyes et al., 1995; Predel et al., 2004; Wegener et al., 2006). When the APCs are activated by stimuli such as peptide hormones or neurons that make physical connections with the APCs (Kubrak et al., 2020; Oh et al., 2019; Scopelliti et al., 2019; Zhao and Karpac, 2017), Akh is released into the hemolymph (Braco et al., 2012).

Circulating Akh then interacts with a G-protein coupled receptor called the Akh receptor (AkhR, FBgn0025595), where Akh binding to AkhR on target tissues such as the fat body increases intracellular cyclic adenosine monophosphate (cAMP) levels.

High levels of cAMP activate protein kinase A (PKA; FBgg0000242) (Gäde and Auerswald, 2003; Park et al., 2002; Staubli et al., 2002a), which phosphorylates several downstream metabolic effectors to promote fat breakdown. For example, in insects, active PKA promotes fat breakdown via phosphorylation and activation of Lipid storage droplet-1 (Lsd-1; FBgn0039114) (Arrese et al., 2008; Bickel et al., 2009; Gäde and Auerswald, 2003; Patel et al., 2005). In mammals, fat breakdown is mediated by similar PKA-dependent phosphorylation of mammalian Lsd-1, and by PKA-dependent phosphorylation and recruitment of lipases, such as Hormone-sensitive lipase (Hsl), to lipid droplets to promote fat mobilization (Sztalryd and Brasaemle, 2017). Given that these genes are highly conserved in flies (Kühnlein, 2012), similar PKA-dependent mechanisms likely explain triglyceride mobilization from lipid droplets. Thus, high levels of Akh pathway activity limit fat storage whereas low levels of Akh signaling promote fat storage. While Akh-mediated triglyceride breakdown plays a vital role in releasing stored energy during times of nutrient scarcity to promote survival (Mochanová et al., 2018), the Akh pathway limits fat storage even in contexts when nutrients are plentiful. Indeed, loss of *Akh* or *AkhR* augments fat storage in males under normal physiological conditions (Bharucha et al., 2008; Gáliková et al., 2015; Grönke et al., 2007), highlighting the critical role of this pathway in regulating whole-body triglyceride levels.

Additional clues into potential mechanisms underlying the sex difference in fat storage come from studies on metabolic genes. For example, flies carrying loss-of-function mutations in genes involved in triglyceride synthesis and storage, such as *midway* (*mdy*; FBgn0004797), *Lipin* (*Lpin;* FBgn0263593), *Lipid storage droplet-2* (*Lsd- 2*; FBgn0030608), and *Seipin* (*Seipin*; FBgn0040336) show reduced whole-body triglyceride levels (Buszczak et al., 2002; Grönke et al., 2003; Teixeira et al., 2003; Tian et al., 2011; Ugrankar et al., 2011; Wang et al., 2016). Whole-body deficiency for genes that regulate triglyceride breakdown, on the other hand, generally have higher whole-body fat levels. This is best illustrated by elevated whole-body triglyceride levels found in flies lacking *brummer* (*bmm*; FBgn0036449) or *Hsl* (FBgn0034491), both of which encode lipases (Bi et al., 2012; Grönke et al., 2005). While these studies demonstrate the strength of *Drosophila* as a model in revealing conserved mechanisms that contribute to whole-body fat storage (Recazens et al., 2021; Schreiber et al., 2019; Walther and Farese, 2012), studies on *Drosophila* fat metabolism often use single- or mixed-sex groups of flies (Bednářová et al., 2018; Gáliková et al., 2015; Grönke et al., 2007; Hughson et al., 2021; Isabel et al., 2005; Lee and Park, 2004; Scopelliti et al., 2019). As a result, less is known about how these metabolic genes and pathways contribute to the sex difference in fat storage.

Recent studies have begun to fill this knowledge gap by studying fat metabolism in both sexes. In one study, higher circulating levels of steroid hormone ecdysone in mated females were found to promote increased whole-body fat storage (Sieber and Spradling, 2015). Another study showed that elevated levels of *bmm* mRNA in male flies restricted triglyceride storage to limit whole-body fat storage (Wat et al., 2020). Yet, neither ecdysone signaling nor *bmm* fully explain the male-female differences in whole-body fat metabolism (Sieber and Spradling, 2015; Wat et al., 2020), suggesting that additional metabolic genes and pathways must contribute to the sex difference in fat storage (Wat et al., 2020). Indeed, genome-wide association studies in *Drosophila* support sex-biased effects on fat storage for many genetic loci (Nelson et al., 2016; Watanabe and Riddle, 2021). As evidence of sex-specific mechanisms underlying whole-body fat storage continues to mount, several reports have also identified male-female differences in phenotypes linked with fat metabolism. For example, sex differences have been reported in energy physiology, metabolic rate, food intake, food preference, circadian rhythm, sleep, immune response, starvation resistance, and lifespan (Andretic and Shaw, 2005; Austad and Fischer, 2016; Belmonte et al., 2020; Chandegra et al., 2017; Helfrich-Förster, 2000; Huber et al., 2004; Hudry et al., 2019; Millington et al., 2021; Park et al., 2018; Reddiex et al., 2013; Regan et al., 2016; Sieber and Spradling, 2015; Videlier et al., 2019; Wat et al., 2020). More work is therefore needed to understand the genetic and physiological mechanisms underlying the male-female difference in fat storage, and to identify the impact of this sex-specific regulation on key life history traits. Further, it will be critical to elucidate how these mechanisms are linked with upstream factors that determine sex.

In *Drosophila*, sexual development is determined by the number of X chromosomes (Salz and Erickson, 2010). In females, the presence of two X chromosomes triggers the production of a functional splicing factor called Sex lethal (Sxl; FBgn0264270) (Bell et al., 1988; Bridges, 1921; Cline, 1978). Sxl’s most well-known downstream target is *transformer* (*tra*; FBgn0003741), where Sxl-dependent splicing of *tra* pre-mRNA allows the production of a functional Tra protein (Belote et al., 1989; Boggs et al., 1987; Inoue et al., 1990; Sosnowski et al., 1989). In males, which have only one X chromosome, no functional Sxl or Tra proteins are made (Cline and Meyer, 1996; Salz and Erickson, 2010). Over several decades, a large body of evidence has accumulated showing that Sxl and Tra direct most aspects of female sexual identity, including effects on abdominal pigmentation, egg-laying, neural circuits, and behaviour (Anand et al., 2001; Baker et al., 2001; Billeter et al., 2006; Brown and King, 1961; Burtis and Baker, 1989; Camara et al., 2008; Christiansen et al., 2002; Cline, 1978; Cline and Meyer, 1996; Clough et al., 2014; Dauwalder, 2011; Demir and Dickson, 2005; Goodwin et al., 2000; Hall, 1994; Heinrichs et al., 1998; Hoshijima et al., 1991; Inoue et al., 1992; Ito et al., 1996; Nagoshi et al., 1988; Neville et al., 2014; Nojima et al., 2014; Pavlou et al., 2016; von Philipsborn et al., 2014; Pomatto et al., 2017; Rezával et al., 2014, 2016; Rideout et al., 2007, 2010; Ryner et al., 1996; Sturtevant, 1945). More recently, studies have extended our knowledge of how Sxl and Tra regulate additional aspects of development and physiology such as body size and intestinal stem cell proliferation (Ahmed et al., 2020; Hudry et al., 2016; Millington and Rideout, 2018; Millington et al., 2021; Regan et al., 2016; Rideout et al., 2015; Sawala and Gould, 2017). Yet, the effects of sex determination genes on whole-body fat metabolism remain unknown, indicating a need for more knowledge of how factors that determine sexual identity influence this important aspect of physiology.

Here, we reveal a role for sex determination gene *tra* in regulating whole-body triglyceride storage. In females, Tra expression promotes a higher level of whole-body fat storage, whereas lack of a functional Tra protein in males leads to lower fat storage. Interestingly, neurons were the anatomical focus of *tra*’s effects on fat storage, where we show that ectopic Tra expression in male APCs was sufficient to augment whole-body triglyceride levels. Our analysis of Akh pathway regulation in both sexes revealed increased *Akh/AkhR* mRNA levels, APC activity, and Akh pathway activity in males. Our findings indicate that this overall male bias in the Akh pathway contributes to the sex difference in whole-body triglyceride levels by restricting fat storage in males.

Importantly, we show that the presence of Tra influences Akh pathway activity, and that Akh lies genetically downstream of Tra in regulating whole-body fat storage. These results provide new insight into the mechanisms by which upstream determinants of sexual identity, such as *tra,* influence the sex difference in fat storage. Further, we identify a previously unrecognized sex-biased role for Akh in regulating whole-body triglyceride levels.

## RESULTS

### Sex determination gene *transformer* regulates the male-female difference in fat storage

Altered *Sxl* function in either sex causes significant lethality due to effects on the dosage compensation machinery (Cline, 1978; Cline and Meyer, 1996). We therefore asked whether the presence of Tra in females, which promotes female sexual development, contributes to the elevated whole-body triglyceride levels observed in females (Sieber and Spradling, 2015; Wat et al., 2020). In 5-day-old virgin females lacking *tra* function (*tra^1^/Df(3L)st-j7*), we found that whole-body triglyceride levels were significantly lower than in age-matched *w^1118^* control females (Figure 1A). Because we observed no significant difference in fat storage between *tra^1^/Df(3L)st-j7* mutant males and *w^1118^* controls (Figure 1 - figure supplement 1A), the sex difference in whole-body triglyceride storage was reduced. Importantly, Tra’s effect on whole-body triglyceride storage was not explained by the absence of ovaries in females lacking Tra function (Sieber and Spradling, 2015; Wat et al., 2020), as whole-body fat storage was significantly reduced in *tra^1^/Df(3L)st-j7* mutant females without gonads compared with *w^1118^* control females lacking ovaries (Figure 1B). Given that we reproduced this finding in females carrying a distinct combination of *tra* mutant alleles (Figure 1C) (Hudry et al., 2016), our findings suggest Tra regulates the sex difference in whole-body triglyceride levels by promoting fat storage in females. While females have reduced fat breakdown post-starvation compared with males (Wat et al., 2020), the magnitude of fat breakdown post-starvation was not significantly different between *tra^1^/Df(3L)st-j7* mutants and sex- matched *w^1118^* controls (genotype:time interactions *p*=0.6298 [females], *p*=0.3853 [males]; Supplementary file 1) (Figure 1 - figure supplement 1B). Tra function is therefore required to promote elevated fat storage in females, but does not regulate fat breakdown post-starvation.

**Figure 1.**
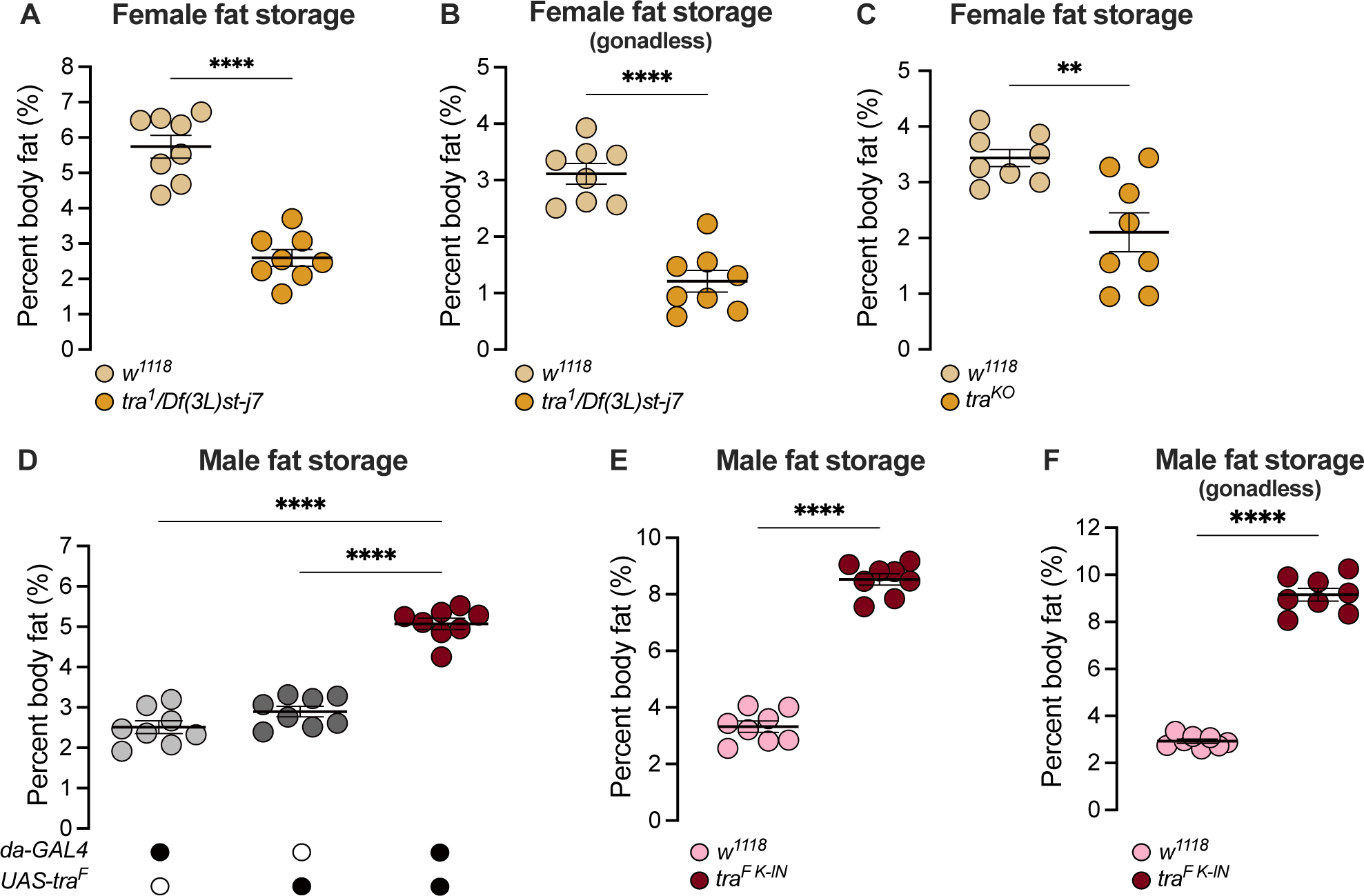
*transformer* regulates the sex difference in fat storage. (A) Whole-body triglyceride levels were significantly lower in *tra^1^/Df(3L)st-j7* females compared with *w^1118^* controls (*p<0.0001*; Student’s *t*-test). *n*=8 biological replicates. (B) Whole-body triglyceride levels were significantly lower in *tra^1^/Df(3L)st-j7* females with excised gonads compared with *w^1118^* controls lacking gonads (*p<0.0001*; Student’s *t*-test). *n*=8 biological replicates. (C) Whole-body triglyceride levels were significantly lower in *tra^KO^* females compared with *w^1118^* controls (*p*=0.0037; Student’s *t*-test). *n*=8 biological replicates. (D) Whole-body triglyceride levels were significantly higher in *da- GAL4>UAS-tra^F^* males compared with *da-GAL4>+* and *+>UAS-tra^F^* controls (*p*<0.0001 and *p*<0.0001 respectively; one-way ANOVA followed by Tukey’s HSD). *n*=8 biological replicates. (E) Whole-body triglyceride levels were significantly higher in *tra^F K-IN^* males compared with *w^1118^* controls (*p*<0.0001, Student’s *t*-test). *n*=8 biological replicates. (F) Whole-body triglyceride levels were significantly higher in *tra^F K-IN^* males with excised gonads compared with *w^1118^* controls lacking gonads (*p<*0.0001; Student’s *t*-test). *n*=8 biological replicates. Black circles indicate the presence of a transgene and open circles indicate the lack of a transgene. ** indicates *p*<0.01, **** indicates *p*<0.0001; error bars represent SEM.

Given that males normally lack a functional Tra protein (Belote et al., 1989; Boggs et al., 1987; Inoue et al., 1990; Sosnowski et al., 1989), we next asked whether the absence of Tra in males explains their reduced whole-body triglyceride levels and rapid triglyceride breakdown post-starvation (Wat et al., 2020). To test this, we ubiquitously overexpressed Tra using *daughterless* (*da*)-*GAL4*, an established way to feminize male flies (Ferveur et al., 1995; Rideout et al., 2015), and examined whole-body fat metabolism. In 5-day-old *da-GAL4>UAS-tra^F^* males, whole-body triglyceride levels were significantly higher than in age-matched *da-GAL4>+* or *+>UAS-tra^F^* control males (Figure 1D). No increase in whole-body fat storage was observed in age-matched *da-GAL4>UAS-tra^F^* females compared with *da-GAL4>+* or *+>UAS-tra^F^* control females (Figure 1 - figure supplement 1C); therefore, the sex difference in fat storage was reduced. Because high levels of Tra overexpression may influence viability (Siera and Cline, 2008), we also measured fat storage in males carrying an allele of *tra* that directs the production of physiological Tra levels (*tra^F K-IN^* allele) (Hudry et al., 2019). As in *da- GAL4>UAS-tra^F^* males, whole-body triglyceride levels were significantly higher in *tra^F K- IN^* males compared with *w^1118^* control males (Figure 1E), indicating that the gain of a functional Tra protein in males promotes elevated whole-body fat storage.

Importantly, the presence of rudimentary ovaries in *tra^F K-IN^* males did not explain their increased fat storage, as whole-body fat storage was still higher in *tra^F K-IN^* males lacking gonads compared with gonadless control males (Figure 1F). The elevated fat storage in *tra^F K-IN^* males also cannot be attributed to ecdysone production by the rudimentary ovaries, as no ecdysone target genes were upregulated (Figure 1 - figure supplement 1D) (Sieber and Spradling, 2015). Together, these data indicate that lack of Tra function contributes to the reduced whole-body triglyceride levels normally observed in males. In males, this role for Tra may also extend to regulation of fat breakdown, as triglyceride mobilization post-starvation was significantly reduced in *da-GAL4>UAS-tra^F^* males compared with *da-GAL4>+* or *+>UAS-tra^F^* controls during a 24 hr starvation period (genotype:time *p*<0.0001 [males]; Supplementary file 1) (Figure 1 - figure supplement 1E), a finding we reproduced in *tra^F K-IN^* males (Figure 1 - figure supplement 1F). While this effect of Tra on fat breakdown in males does not perfectly align with our data from *tra* mutant females, we note a trend toward increased fat breakdown in *tra* mutant females that was not statistically significant (Figure 1 - figure supplement 1B).

Taken together, these data support a clear role for Tra in regulating the sex difference in fat storage, and suggest that a role for Tra in regulating fat breakdown cannot be ruled out.

### *transformer* function in neurons regulates the sex difference in fat storage

Tra function is required in many cell types, tissues, and organs to promote female sexual development (Anand et al., 2001; Baker et al., 2001; Billeter et al., 2006; Brown and King, 1961; Burtis and Baker, 1989; Camara et al., 2008; Christiansen et al., 2002; Clough et al., 2014; Dauwalder, 2011; Demir and Dickson, 2005; Goodwin et al., 2000; Hall, 1994; Heinrichs et al., 1998; Hoshijima et al., 1991; Inoue et al., 1992; Ito et al., 1996; Nagoshi et al., 1988; Neville et al., 2014; Nojima et al., 2014; Pavlou et al., 2016; von Philipsborn et al., 2014; Pomatto et al., 2017; Rezával et al., 2014, 2016; Rideout et al., 2007, 2010; Ryner et al., 1996; Sturtevant, 1945). To determine the cell types and tissues in which Tra function is required to influence fat metabolism, we overexpressed Tra using a panel of GAL4 lines that drive expression in subsets of cells and/or tissues. To rapidly assess potential effects on fat metabolism, we measured starvation resistance, an established readout for changes to fat storage and breakdown (Beller et al., 2010; Bi et al., 2012; Choi et al., 2015; Grönke et al., 2003, 2005, 2007; Gutierrez et al., 2007).

Normally, adult females have elevated starvation resistance compared with age-matched males due to higher fat storage and reduced fat breakdown (Wat et al., 2020). Indeed, loss of *tra* reduced starvation resistance in females (Figure 2A) whereas gain of Tra function enhanced starvation resistance in males (Figure 2B), in line with their effects on fat metabolism (Figure 1A,D). From our survey of different GAL4 lines (Figure 2 - figure supplement 1A-F; Figure 2 - figure supplement 2A-D), we found that neurons were the cell type in which gain of Tra most strongly extended male starvation resistance (Figure 2C). Specifically, starvation resistance in males with Tra overexpression in neurons (*elav-GAL4>UAS-tra^F^*) was significantly extended compared with *elav-GAL4>+* and *+>UAS-tra^F^* controls (Figure 2C), with no effect in females (Figure 2 - figure supplement 3A). Because the increase in starvation resistance upon neuron-specific Tra expression was similar in magnitude to the increase in survival observed upon global Tra expression (Figure 2B,C), this finding suggests a key role for neuronal Tra in regulating starvation resistance.

**Figure 2.**
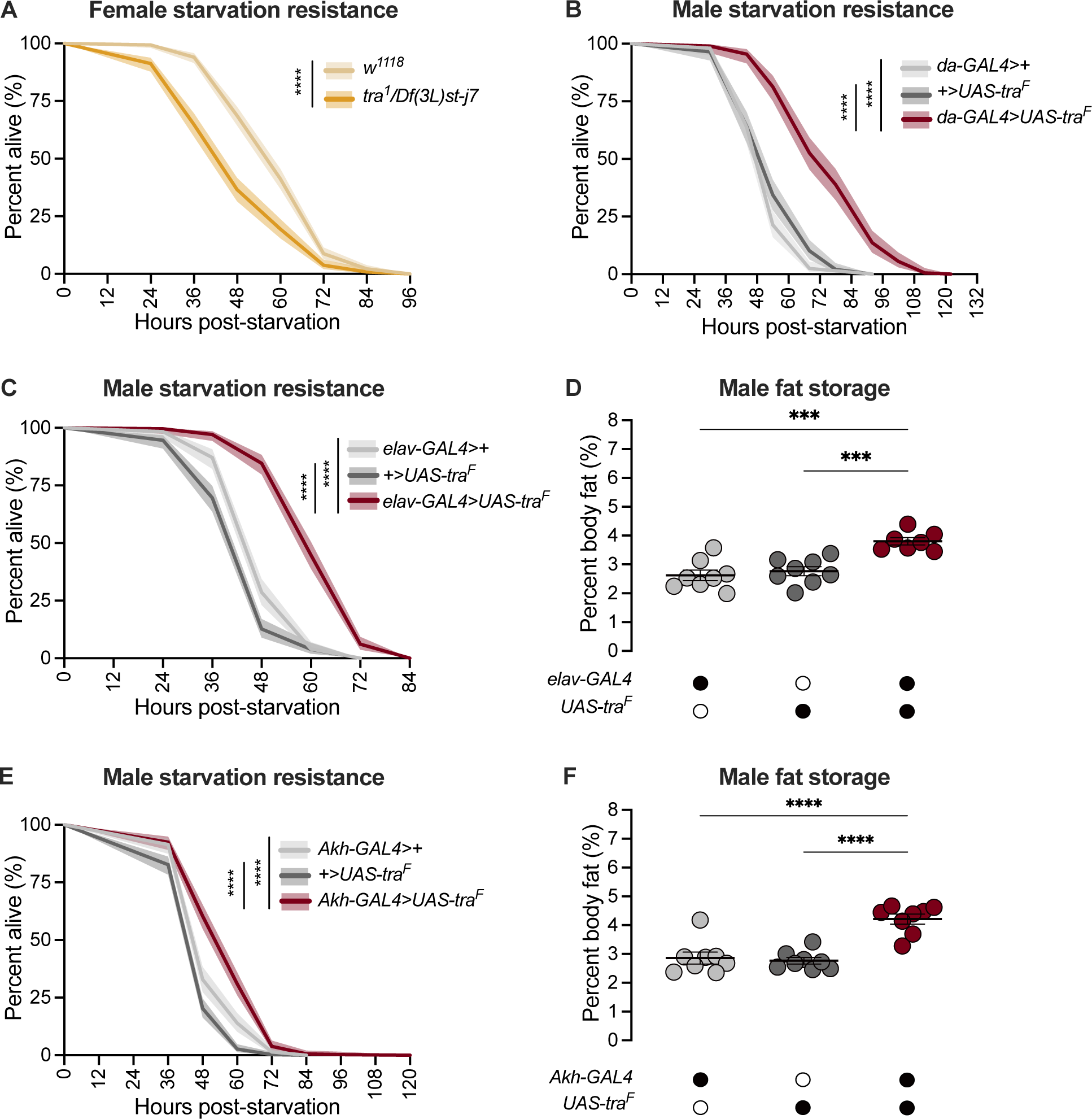
*transformer* function in Akh-producing cells contributes to the sex difference in fat storage. (A) Starvation resistance was significantly reduced in *tra^1^/Df(3L)st-j7* females compared with *w^1118^* controls (*p*<2x10^-16^; log-rank test, Bonferroni’s correction for multiple comparisons). *n*=344-502 animals. (B) Starvation resistance was significantly enhanced in *da-GAL4>UAS-tra^F^* males compared with *da- GAL4>+* and *+>UAS-tra^F^* controls (*p*<2x10^-16^ and *p*<2x10^-16^ respectively; log-rank test, Bonferroni’s correction for multiple comparisons). *n*=198-201 animals. (C) Starvation resistance was significantly enhanced in *elav-GAL4>UAS-tra^F^* males compared with *elav-GAL4>+* and *+>UAS-tra^F^* controls (*p*<2x10^-16^ and *p*<2x10^-16^ respectively; log-rank test, Bonferroni’s correction for multiple comparisons). *n*=248-279 animals. (D) Whole- body triglyceride levels were significantly higher in *elav-GAL4>UAS-tra^F^* males compared with *elav-GAL4>+* and *+>UAS-tra^F^* controls (*p*=0.0001 and *p*=0.0006 respectively; one-way ANOVA followed by Tukey’s HSD). *n*=7-8 biological replicates. (E) Starvation resistance was significantly enhanced in *Akh-GAL4>UAS-tra^F^* males compared with *Akh-GAL4>+* and *+>UAS-tra^F^* controls (*p*=3.1x10^-9^ and *p*<2x10^-16^ respectively; log-rank test, Bonferroni’s correction for multiple comparisons). *n*=280-364 animals. (F) Whole-body triglyceride levels were significantly higher in *Akh-GAL4>UAS- tra^F^* males compared to *Akh-GAL4>+* and *+>UAS-tra^F^* control males (*p*<0.0001 and *p*<0.0001 respectively; one-way ANOVA followed by Tukey’s HSD). *n*=8 biological replicates. Black circles indicate the presence of a transgene and open circles indicate the lack of a transgene. *** indicates *p*<0.001, **** indicates *p*<0.0001; shaded areas represent the 95% confidence interval; error bars represent SEM.

To determine whether increased starvation resistance in *elav-GAL4>UAS-tra^F^* males was due to altered fat metabolism, we measured whole-body triglyceride levels in males and females with neuronal Tra overexpression. We found that *elav-GAL4>UAS- tra^F^* males (Figure 2D), but not females (Figure 2 - figure supplement 3B), showed a significant increase in whole-body fat storage compared with sex-matched *elav-GAL4>+* and *+>UAS-tra^F^* controls. This suggests that the male-specific increase in starvation resistance (Figure 2C) was due to increased fat storage in *elav-GAL4>UAS-tra^F^* males, which we confirm by showing that the rate of fat breakdown in *elav-GAL4>UAS-tra^F^* males and females was not significantly different from sex-matched *elav-GAL4>+* and *+>UAS-tra^F^* controls (Figure 2 - figure supplement 3C) (genotype:time interaction *p*=0.2789 [males], *p*=0.7058 [females]; Supplementary file 1). Neurons are therefore one cell type in which Tra function influences the sex difference in whole-body triglyceride storage.

To identify specific neurons that mediate Tra’s effects on starvation resistance and whole-body fat storage, we overexpressed Tra in neurons known to affect fat metabolism and measured starvation resistance (Figure 2 - figure supplement 4A-E) (Al-Anzi and Zinn, 2018; Al-Anzi et al., 2009; Chung et al., 2017; Li et al., 2016; May et al., 2020; Min et al., 2016; Mosher et al., 2015; Zhan et al., 2016). One group of neurons that significantly augmented starvation resistance upon Tra expression was the APCs (Figure 2E), a group of neuroendocrine cells in the corpora cardiaca that produce Akh and other peptide hormones such as Limostatin (Lst; FBgn0034140) (Alfa et al., 2015; Lee and Park, 2004). Flies with APC-specific Tra expression (*Akh-GAL4>UAS- tra^F^)* had significantly increased starvation resistance compared with sex-matched *Akh- GAL4>+* and *+>UAS-tra^F^* controls (Figure 2E; Figure 2 - figure supplement 5A). To determine whether the starvation resistance phenotype indicated altered fat storage, we compared whole-body triglyceride levels in *Akh-GAL4>UAS-tra^F^* males and females with sex-matched *Akh-GAL4>+* and *+>UAS-tra^F^* controls. There was a significant increase in whole-body fat storage in males (Figure 2F) but not females (Figure 2 - figure supplement 5B) with APC-specific Tra expression. This indicates Tra function in the APCs promotes fat storage, revealing a previously unrecognized role for the APCs in regulating the sex difference in fat storage. Indeed, fat breakdown was unaffected in *Akh-GAL4>UAS-tra^F^* males and females compared with sex-matched *Akh-GAL4>+* and *+>UAS-tra^F^* controls (Figure 2 - figure supplement 5C) (genotype:time interaction *p*=0.1201 [males] and *p*=0.0596 [females]; Supplementary file 1).

### Sex-specific regulation of Adipokinetic hormone leads to a male bias in pathway activity

Given that the sexual identity of the APCs impacts whole-body fat storage, we compared the regulation of *Akh*, APC activity, and Akh signaling between adult males and females. We first examined *Akh* and *AkhR* mRNA levels in both sexes using quantitative real-time polymerase chain reaction (qPCR). We found that mRNA levels of both *Akh* and *AkhR* were significantly higher in 5-day-old *w^1118^* males than in females (Figure 3A,B). This male bias in *Akh* mRNA levels did not reflect an increased APC number in males, as we found no sex difference in the number of APCs (Figure 3C).

**Figure 3.**
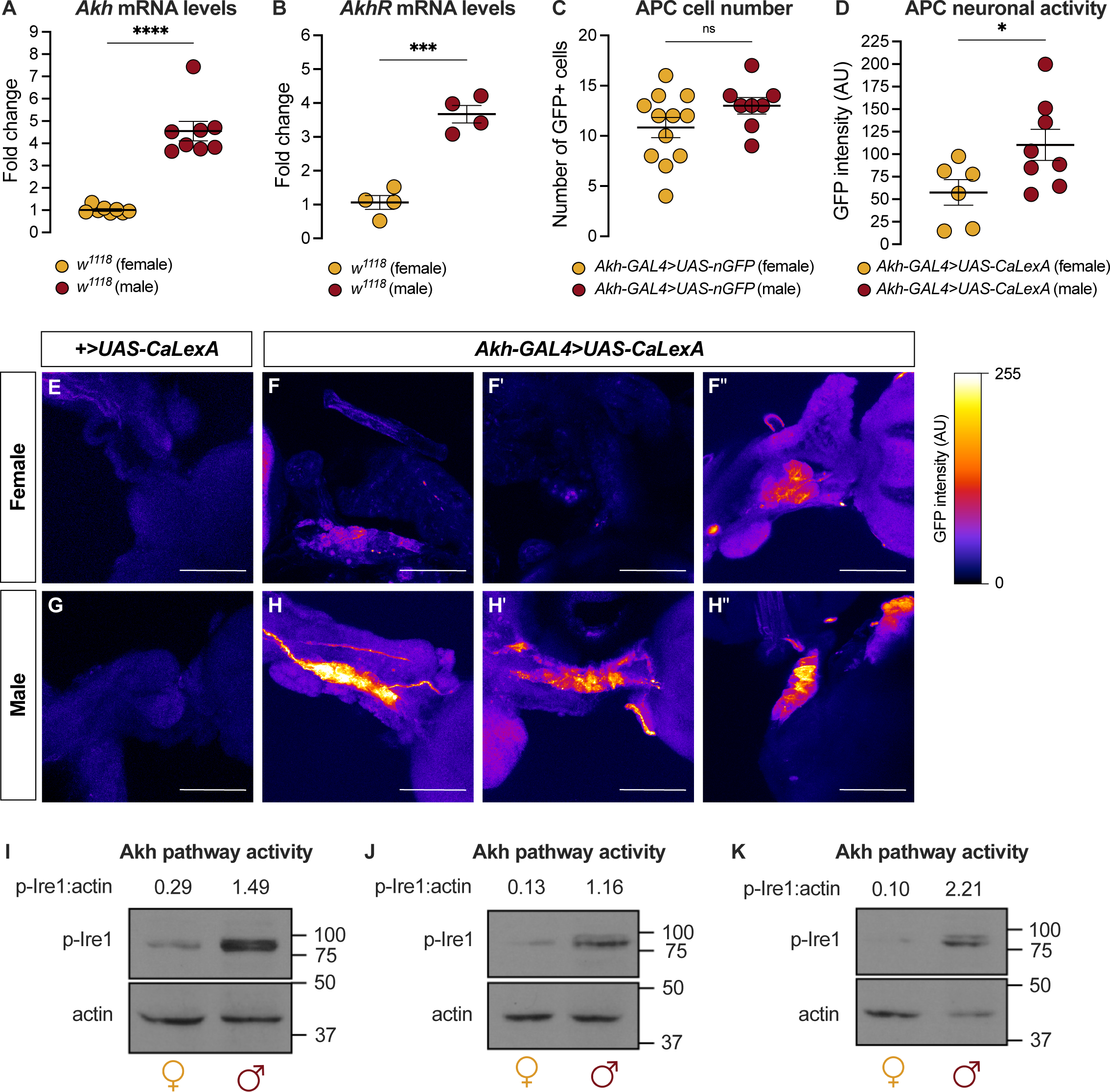
Sex-specific regulation of Akh and the Akh signaling pathway. (A) *Akh* mRNA levels were significantly higher in *w^1118^* males compared with genotype-matched females (*p<*0.0001, Student’s *t*-test). *n*=8 biological replicates. (B) *AkhR* mRNA levels were significantly higher in *w^1118^* males than in females (*p=*0.0002, Student’s *t*-test). *n*=4 biological replicates. (C) Expression of *UAS-nGFP* in Akh-producing cells (APCs) (*Akh- GAL4>UAS-nGFP*) revealed no significant difference in APC cell number between males and females (*p*=0.1417; Student’s *t*-test). *n*=8-12 animals. (D) GFP intensity produced as a readout of calcium activity in the APCs (*Akh-GAL4>LexAop-CD8- GFP;UAS-LexA-VP16-NFAT (UAS-CaLexA)*) was significantly higher in males compared with females (*p*=0.0438; Student’s *t*-test). *n*=6-8 biological replicates. (E-H) Maximum Z-projections of representative images showing GFP produced as a readout for APC calcium activity from both *Akh-GAL4>UAS-CaLexA* males and females. Scale bars=50 μm, *n*=6-8 biological replicates. (I-K) Whole-body p-Ire1 levels were higher in *w^1118^* males compared with *w^1118^* females in three biological replicates. * indicates *p*<0.05, *** indicates *p*<0.001, **** indicates *p*<0.0001, ns indicates not significant; error bars represent SEM. Original images for Figure 3C are found in Figure 3 – Source Data 1. Original images for Figure 3D-H are found in Figure 3 – Source Data 2. Original images for Figure 3I-K are found in Figure 3 – Source Data 3.

Because Akh release from the APCs is regulated by APC activity (Kubrak et al., 2020; Oh et al., 2019), we next measured APC activity in males and females by driving APC-specific expression of calcium-responsive chimeric transcription factor *LexA-VP16- NFAT* (*Akh-GAL4>UAS-LexA-VP16-NFAT* [called *UAS-CaLexA*]) (Masuyama et al., 2012). Sustained APC activity triggers nuclear import of LexA-VP16-NFAT, where it drives expression of a GFP reporter downstream of a LexA-responsive element (Masuyama et al., 2012). Monitoring GFP levels in the APCs therefore provides a straightforward way to monitor APC activity.

In 5-day-old *Akh-GAL4*>*UAS-CaLexA* males, GFP levels were significantly higher than in age- and genotype-matched females (Figure 3D-H). Because *GAL4* mRNA levels were not significantly different between males and females carrying the *Akh-GAL4* transgene (Figure 3 - figure supplement 1A), and the number of APCs did not differ between the sexes (Figure 3C), these findings indicate that the APCs are more active in males than in females. To determine whether the male bias in *Akh/AkhR* mRNA levels and APC activity affected Akh pathway activity, we next compared levels of phosphorylated Inositol-requiring enzyme-1 (Ire1; FBgn0261984) between males and females. Because levels of phosphorylated Ire1 (p-Ire1) are higher in *Drosophila* cells stimulated with Akh peptide, regulation that was dependent on the presence of AkhR, high p-Ire1 levels indicate increased Akh pathway activity (Song et al., 2017). We found that the ratio of p-Ire1 to loading control actin was higher in 5-day-old *w^1118^* males compared with age- and genotype-matched females in three out of four biological replicates (Figure 3I-K; Figure 3 - figure supplement 1B), a finding that aligns with the sex difference in *Akh/AkhR* mRNA levels and APC activity. Taken together, our data suggests a previously unrecognized male bias in the Akh pathway.

### The Adipokinetic hormone pathway contributes to the sex difference in fat storage

Given that high Akh pathway activity limits fat storage via an established intracellular signaling cascade that culminates in lipase recruitment and fat mobilization (Baumbach et al., 2014; Grönke et al., 2007; Lee and Park, 2004; Mochanová et al., 2018), we wanted to determine whether the male bias in Akh pathway activity influences the sex difference in fat metabolism by restricting fat storage in males. We therefore used a published approach to ablate the APCs (*Akh-GAL4>UAS-reaper* (*rpr*)) (Lee and Park, 2004; White et al., 1996), and measured whole-body triglyceride levels in each sex.

Because the sexual identity of the APCs affects fat storage and not fat breakdown (Figure 2F; Figure 2 - figure supplement 5C), we focused our analysis on triglyceride storage rather than mobilization. Triglyceride levels were significantly higher in 5-day-old *Akh-GAL4>UAS-rpr* males than in *Akh-GAL4>+* and *+>UAS-rpr* control males (Figure 4A). In contrast, triglyceride levels in 5-day-old *Akh-GAL4>UAS-rpr* females were not significantly different from *Akh-GAL4>+* and *+>UAS-rpr* control females (Figure 4 - figure supplement 1A). This suggests that the male bias in Akh pathway activity normally contributes to the sex difference in fat storage by limiting triglyceride accumulation in males via the established intracellular signaling cascade known to regulate lipid droplet breakdown (Arrese et al., 2008; Heier and Kühnlein, 2018; Heier et al., 2021; Kühnlein, 2012; Patel et al., 2005). Importantly, we reproduced the male-biased effects on fat storage in flies carrying mutant *Akh* and *AkhR* alleles (*Akh^A^* and *AkhR^1^,* respectively) (Figure 4B,C; Figure 4 - figure supplement 1B,C), and show that APC-specific knockdown of *Lst* had no effect on fat storage in either sex (Figure 4 - figure supplement 1D,E). These findings support a model in which it is Akh production by the APCs that plays a role in regulating the male-female difference in fat storage.

**Figure 4.**
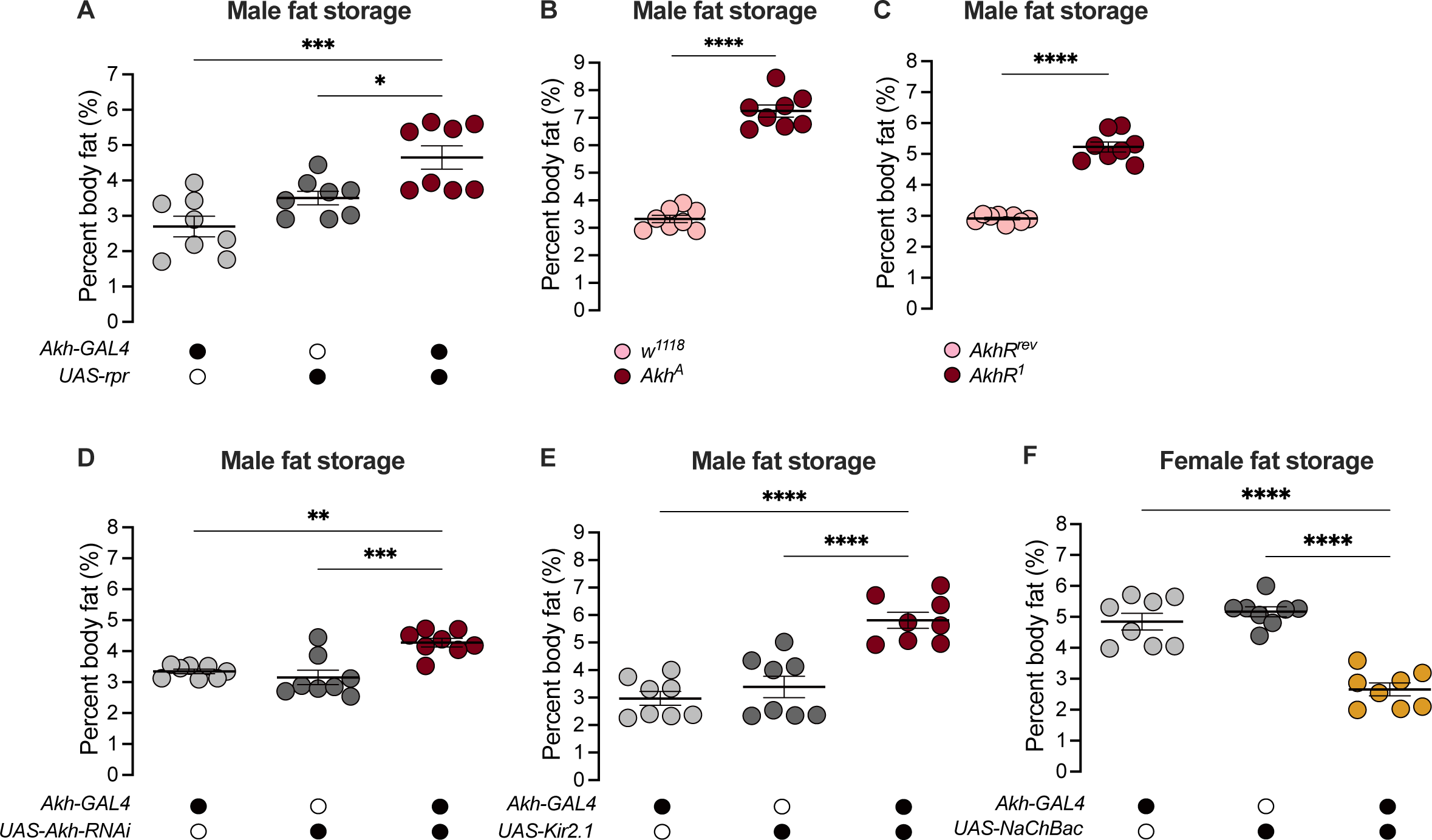
Sex-specific regulation of Akh and APC activity influence the sex difference in fat storage. (A) Whole-body triglyceride levels were significantly higher in *Akh-GAL4>UAS-reaper (rpr)* males compared with *Akh-GAL4>+* and *+>UAS-rpr* controls (*p*=0.0002 and *p*=0.0215 respectively; one-way ANOVA followed by Tukey’s HSD). *n*=8 biological replicates. (B) Whole-body triglyceride levels were significantly higher in *Akh^A^* males compared with *w^1118^* controls (*p*<0.0001; one-way ANOVA followed by Tukey’s HSD). *n*=8 biological replicates. (C) Whole-body triglyceride levels were significantly higher in *AkhR^1^* males compared with *AkhR^rev^* controls (*p*<0.0001; one-way ANOVA followed by Tukey’s HSD). *n*=8 biological replicates. (D) Whole-body triglyceride levels were significantly higher in *Akh-GAL4>UAS-Akh-RNAi* males compared with *Akh-GAL4>+* and *+>UAS-Akh-RNAi* controls (*p*=0.0015 and *p*=0.0002 respectively; one-way ANOVA followed by Tukey’s HSD). *n*=8 biological replicates. (E) Whole-body triglyceride levels were significantly higher in *Akh-GAL4>UAS-Kir2.1* males compared with *Akh-GAL4>+* and *+>UAS-Kir2.1* controls (*p*<0.0001 and *p*<0.0001 respectively; one-way ANOVA followed by Tukey’s HSD). *n*=8 biological replicates. (F) Whole-body triglyceride levels were significantly lower in *Akh-GAL4>UAS-NaChBac* females compared with *Akh-GAL4>+* and *+>UAS-NaChBac* controls (*p*<0.0001 and *p*<0.0001 respectively; one-way ANOVA followed by Tukey’s HSD). *n*=8 biological replicates. Due to independent experiments with a shared GAL4 control, *Akh-GAL4>+* males are shared between Figure 4E and Figure 4 – figure supplement 3E. *Akh- GAL4>+* females are shared between Figure 4F and Figure 4 – figure supplement 3D. Black circles indicate the presence of a transgene and open circles indicate the lack of a transgene; * indicates *p*<0.05, ** indicates *p*<0.01, *** indicates *p*<0.001, **** indicates *p*<0.0001; error bars represent SEM.

While the mechanisms underlying the regulation of intracellular fat breakdown by Akh have been well-documented (Bharucha et al., 2008; Gáliková et al., 2015; Grönke et al., 2007; Heier and Kühnlein, 2018; Heier et al., 2021; Kubrak et al., 2020; Lee and Park, 2004; Lehmann, 2018; Scopelliti et al., 2019; Zhao and Karpac, 2017), our findings reveal a new role for Akh in regulating the sex difference in fat storage. Notably, this Akh-mediated regulation of the male-female difference in fat storage operates in a parallel pathway to the previously described sex-specific role of triglyceride lipase *bmm* (Figure 4 - figure supplement 2A,B) (Wat et al., 2020).

Beyond the APC ablation or complete loss of Akh, we next wanted to test whether the sex-specific Akh regulation we uncovered contributes to the male-female difference in fat storage. To this end, we used a genetic approach to manipulate *Akh* mRNA levels or APC activity, and measured whole-body fat storage in both sexes. To determine whether the male bias in *Akh* mRNA levels contributes to the sex difference in fat storage, we measured whole-body triglyceride levels in flies with APC-specific expression of *Akh-RNAi* (*Akh-GAL4>UAS-Akh-RNAi*). Importantly, this manipulation effectively reduced *Akh* mRNA levels in both sexes (Figure 4 - figure supplement 3A,B). In males, whole-body triglyceride levels were significantly higher in *Akh-GAL4>UAS-Akh-RNAi* flies compared with *Akh-GAL4>+* and *+>UAS-Akh-RNAi* control flies (Figure 4D). *Akh-GAL4>UAS-Akh-RNAi* female flies, in contrast, showed no significant change in whole-body fat storage compared with *Akh-GAL4>+* and *+>UAS-Akh-RNAi* control females (Figure 4 - figure supplement 3C). This indicates a strongly male-biased effect on fat storage due to reduced *Akh* mRNA levels, suggesting that the sex difference in *Akh* mRNA levels contributes to the male-female difference in whole-body fat storage.

To determine whether the male bias in APC activity also influences the sex difference in fat storage, we silenced the APCs by APC-specific overexpression of an inwardly rectifying potassium channel Kir2.1 (Baines et al., 2001) and measured whole-body triglyceride levels. Whole-body fat storage in *Akh-GAL4>UAS-Kir2.1* adult males was significantly higher compared with *Akh-GAL4>+* and *+>UAS-Kir2.1* control males (Figure 4E). In females, while we observed significantly elevated whole-body fat storage in *Akh-GAL4>UAS-Kir2.1* adults compared with *Akh-GAL4*>+ and +>*UAS-Kir2.1* controls (Figure 4 - figure supplement 3D), the magnitude of this increase was larger in males (sex:genotype interaction *p*=0.0455; Supplementary file 1). Together, these data suggest that the male bias in APC activity contributes to the sex difference in fat storage by limiting triglyceride accumulation in males. Indeed, augmenting APC activity in females using a bacterial voltage-gated sodium channel (*UAS-NaChBac*) significantly reduced fat storage in females (Figure 4F; Figure 4 - figure supplement 3E). While Akh affects food-related behaviours in some contexts (Choi et al., 2015; Hentze et al., 2015; Huang et al., 2020), we observed no significant effects of altered APC activity on feeding behaviour in either sex (Figure 4 - figure supplement 4A-D). This suggests that the male-biased effect of APC manipulation on fat storage cannot be fully explained by effects on food intake. Thus, in addition to the contribution of elevated *Akh* mRNA levels in males to the sex difference in fat storage, we also identify a role for the male bias in APC activity in the sex-specific regulation of whole-body triglyceride levels.

### *transformer* regulates the sex difference in fat storage via the Adipokinetic hormone pathway

Given that Tra function and the Akh pathway both contribute to the male-female difference in fat storage, we asked whether the presence of Tra affects the sex bias in Akh pathway activity. In 5-day-old *tra^1^/Df(3L)st-j7* females, levels of p-Ire1 were higher than in *w^1118^* control females in three out of four biological replicates (Figure 5A-C; Figure 5 - figure supplement 1A). This suggests the presence of Tra in females normally represses Akh pathway activity. Indeed, loss of Tra significantly increased *Akh* mRNA levels in females (Figure 5D). Given Tra’s effects on Akh pathway activity, we next tested whether the change in Akh pathway function was significant for Tra’s effects on whole-body triglyceride levels. We predicted that if increased Akh pathway activity caused the lower fat storage in *tra* mutant females, genetic manipulations that reduce Akh pathway activity should block this reduction in whole-body triglyceride levels. While all female genotypes lacking *tra* function had reduced fat storage compared with control females (Figure 5E), APC ablation in *tra* mutant females rescued this decrease in whole-body triglyceride levels (Figure 5E). Indeed, fat storage in *tra* mutant females lacking APCs was not significantly different from *w^1118^* control females (*p*=0.9384; Supplementary file 1) (Figure 5E), indicating that the increased Akh pathway activity we observed in *tra* mutant females was one reason for their reduced fat storage. Given that APC activation in males expressing physiological levels of Tra similarly rescued the Tra-induced increase in whole-body triglyceride levels (Figure 5F), these findings suggest that the sex-specific regulation of Akh pathway activity represents one way *tra* influences the male-female difference in fat storage.

**Figure 5.**
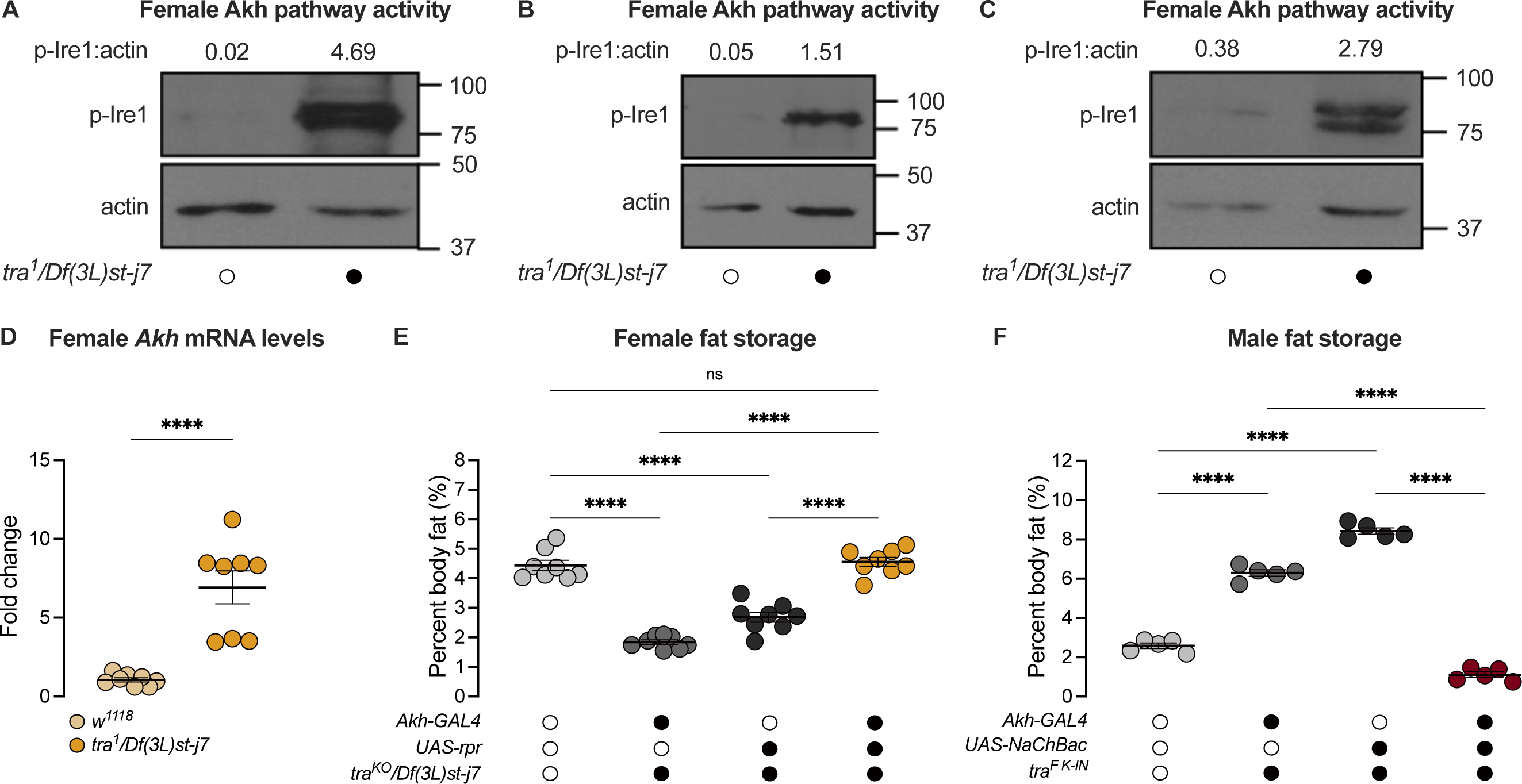
*transformer* regulates the sex difference in fat storage via the Akh signalling pathway. (A-C) Whole-body p-Ire1 levels were higher in *tra^1^/Df(3L)st-j7* females compared with *w^1118^* controls in three biological replicates. (D) Whole-body *Akh* mRNA levels were significantly higher in *tra^1^/Df(3L)st-j7* females compared with *w^1118^* controls (*p*<0.0001; Student’s *t*-test). *n*=8 biological replicates. (E) Whole-body triglyceride levels were significantly lower in *tra^KO^/Df(3L)st-j7* females carrying either *Akh-GAL4>+* or *+>UAS-reaper* (*rpr*) transgenes compared with *w^1118^* controls carrying a functional Tra protein (*p*<0.0001 and *p*<0.0001 respectively; one-way ANOVA followed by Tukey’s HSD). Whole-body triglyceride levels were not significantly different between *tra^KO^/Df(3L)st-j7* females lacking APCs (*Akh-GAL4>UAS-rpr*) and *w^1118^* controls (*p*=0.9384; one-way ANOVA followed by Tukey’s HSD). *n*=8 biological replicates. (F) Whole-body triglyceride levels were significantly higher in *tra^F^* ^K-IN^ males carrying either *Akh-GAL4>+* or *+>UAS-NaChBac* transgenes compared with *w^1118^* control males lacking Tra function (*p*<0.0001 and *p*<0.0001 respectively; one-way ANOVA followed by Tukey’s HSD). Whole-body triglyceride levels in *tra^F^* ^K-IN^ males with APC activation (*Akh- GAL4>UAS-NaChBac*) were significantly lower than *tra^F^* ^K-IN^ males carrying either the *Akh-GAL4>+* or *+>UAS-NaChBac* transgenes alone (*p*<0.0001 and *p*<0.0001 respectively; one-way ANOVA followed by Tukey’s HSD). *n*=5 biological replicates. Black circles indicate the presence of a transgene or mutant allele and open circles indicate the lack of a transgene or mutant allele. **** indicates *p*<0.0001, ns indicates not significant; error bars represent SEM. Original images for Figure 5A-C are found in Figure 5 – Source Data 1.

### Loss of Adipokinetic hormone has opposite effects on reproductive success in each sex and mediates a fecundity-lifespan tradeoff in females

Our results suggest that adult females show lower Akh pathway activity and higher fat storage, whereas males maintain a higher level of Akh activity and lower fat storage. Because the correct regulation of fat storage in each sex influences reproduction (Buszczak et al., 2002; Grönke et al., 2005; Sieber and Spradling, 2015; Wat et al., 2020), we tested how loss of this critical regulator of the sex difference in fat storage impacted offspring production in each sex. In *Akh^A^* mutant males, we found that the proportion of males copulating with a *Canton-S (CS)* virgin female was lower than in control *w^1118^* males at each 10 min interval during a 60 min observation period (Figure 6A). When we counted viable offspring from these copulation events, we found that *Akh^A^* mutant males had significantly fewer overall progeny than *w^1118^* control males (Figure 6B). These results suggest that Akh function normally promotes reproductive success in males; however, it is important to note that Akh function is not absolutely required for male fertility, as a prolonged 24 hr period of contact between *Akh^A^* mutant males and *CS* females allowed the production of normal progeny numbers (Figure 6C).

**Figure 6.**
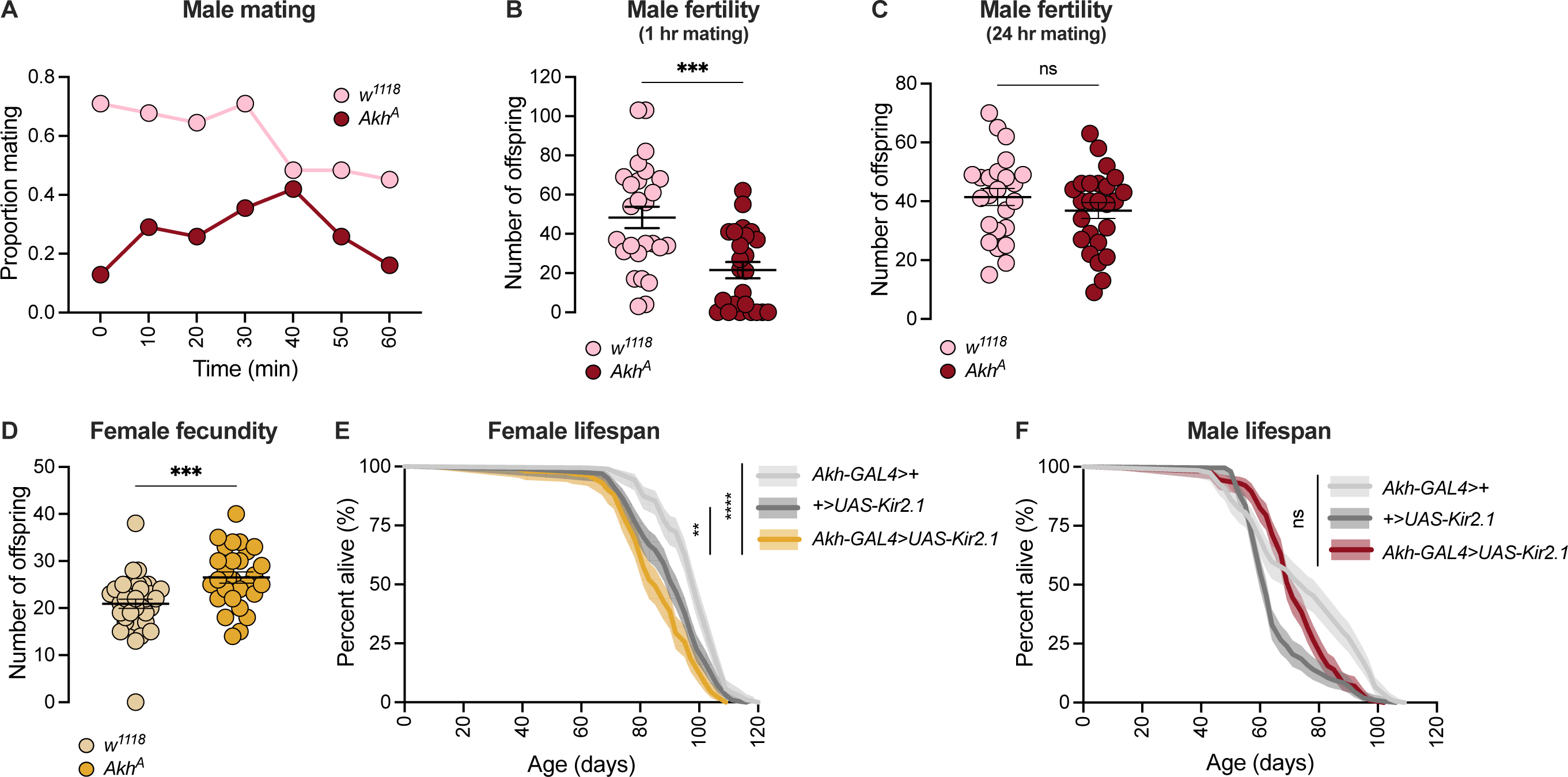
Sex-specific regulation of Akh signalling pathway promotes reproductive success in each sex. (A) At all observation points, a lower proportion of *Akh^A^* males were successfully copulating with a wildtype *Canton-S* (*CS*) female compared with *w^1118^* controls. *n*=31 males. (B) The number of pupae produced from a 60 min mating period was significantly lower in *Akh^A^* males compared with *w^1118^* controls (*p*=0.0003; Student’s *t*-test). *n*=24-26 males. (C) The number of pupae produced from a 24 hr mating period was not significantly different between *Akh^A^* males and *w^1118^* control males (*p*=0.2501; Student’s *t*-test). *n*=24-25 males. (D) The number of pupae produced from a 24 hr mating period was significantly higher in *Akh^A^* females compared with *w^1118^* controls (*p*=0.0006; Student’s *t*-test). *n*=28-36 females. (E) Lifespan was significantly shorter in *Akh-GAL4>UAS-Kir2.1* females compared with *Akh-GAL4>+* and *+>UAS- Kir2.1* controls (*p*<2x10^-16^ and *p*=0.0015 respectively; log-rank test, Bonferroni’s correction for multiple comparisons). *n*=160-198 females. (F) Lifespan of *Akh- GAL4>UAS-Kir2.1* males was intermediate between *Akh-GAL4>+* and *+>UAS-Kir2.1* controls, indicating no overall effect of inhibiting APC neuronal activity on male lifespan (*p*=0.00013 and *p*=7.0x10^-6^ respectively; log-rank test, Bonferroni’s correction for multiple comparisons). *n*=196-200 males. ** indicates *p*<0.01, *** indicates *p*<0.001, **** indicates *p*<0.0001, ns indicates not significant; error bars represent SEM; shaded areas represent the 95% confidence interval.

In contrast to males, loss of Akh in females increased fecundity (Figure 6D).

Specifically, *Akh^A^* mutant females produced a significantly higher number of offspring compared with *w^1118^* controls (Figure 6D). Thus, in females, a low level of Akh pathway activity promotes fecundity. Given that a change in one life history trait such as reproduction often affects traits such as longevity (Chapman et al., 1995; Flatt, 2011; Fowler and Partridge, 1989; Hansen et al., 2013), we also measured lifespan in females with reduced Akh pathway function. We found that lifespan was significantly shorter in *Akh-GAL4>UAS-Kir2.1* females compared with *Akh-GAL4>+* and *+>UAS-Kir2.1* control females (Figure 6E). In contrast, male lifespan was not significantly different between *Akh-GAL4>UAS-Kir2.1* flies and *Akh-GAL4>+* and *+>UAS-Kir2.1* controls (Figure 6F). Our findings are in agreement with a previous study that demonstrated a female-specific reduction in lifespan in response to whole-body loss of *Akh* (Bednářová et al., 2018).

This suggests that while low Akh activity in females promotes fertility, this benefit comes at the cost of a shorter lifespan.

## DISCUSSION

In this study, we used the fruit fly *Drosophila melanogaster* to improve knowledge of the mechanisms underlying the male-female difference in whole-body triglyceride levels.

We show that the presence of a functional Tra protein in females, which directs many aspects of female sexual development, promotes whole-body fat storage. Tra’s ability to promote fat storage arises largely due to its function in neurons, where we identified the APCs as one neuronal population in which Tra function influences whole-body triglyceride levels. Our examination of *Akh*/*AkhR* mRNA levels and APC activity revealed several differences between the sexes, where these differences lead to higher Akh pathway activity in males than in females. Genetic manipulation of APCs and Akh pathway activity suggest a model in which the sex bias in Akh pathway activity contributes to the male-female difference in fat storage by limiting whole-body triglyceride storage in males. Importantly, we show that Tra function influences Akh pathway activity, and that Akh acts genetically downstream of Tra in regulating whole-body triglyceride levels. This reveals a previously unrecognized genetic and physiological mechanism that contributes to the sex difference in fat storage.

One key finding from our study was the identification of sex determination gene *tra* as an upstream regulator of the male-female difference in fat storage. In females, a functional Tra protein promotes fat storage, whereas lack of Tra in males leads to reduced fat storage. While an extensive body of literature has demonstrated important roles for *tra* in regulating neural circuits, behaviour, abdominal pigmentation, and gonad development (Anand et al., 2001; Baker et al., 2001; Billeter et al., 2006; Brown and King, 1961; Burtis and Baker, 1989; Camara et al., 2008; Christiansen et al., 2002; Clough et al., 2014; Dauwalder, 2011; Demir and Dickson, 2005; Goodwin et al., 2000; Hall, 1994; Heinrichs et al., 1998; Hoshijima et al., 1991; Inoue et al., 1992; Ito et al., 1996; Nagoshi et al., 1988; Neville et al., 2014; Nojima et al., 2014; Pavlou et al., 2016; von Philipsborn et al., 2014; Pomatto et al., 2017; Rezával et al., 2014, 2016; Rideout et al., 2007, 2010; Ryner et al., 1996; Sturtevant, 1945), uncovering a role for *tra* in regulating fat storage significantly extends our understanding of how sex differences in metabolism arise. Given that sex differences exist in other aspects of metabolism (*e.g*. oxygen consumption) (Wat et al., 2020), this new insight suggests that more work will be needed to determine whether *tra* contributes to sexual dimorphism in additional metabolic traits. Indeed, one study showed that *tra* influences the sex difference in adaptation to hydrogen peroxide stress (Pomatto et al., 2017). Beyond metabolism, Tra also regulates multiple aspects of development and physiology such as intestinal stem cell proliferation (Ahmed et al., 2020; Hudry et al., 2016; Millington and Rideout, 2018), carbohydrate metabolism (Hudry et al., 2019), body size (Mathews et al., 2017; Rideout et al., 2015), phenotypic plasticity (Millington et al., 2021), and lifespan responses to dietary restriction (Regan et al., 2016). Because some, but not all, of these studies identify a cell type in which Tra function influences these diverse phenotypes, future studies will need to determine which cell types and tissues require Tra expression to establish a female metabolic and physiological state. Indeed, recent single-cell analyses reveal widespread gene expression differences in shared cell types between the sexes (Li et al., 2021).

Identifying neurons as the anatomical focus of Tra’s effects on fat storage was another key finding from our study. While many sexually dimorphic neural circuits related to behaviour and reproduction have been identified (Anand et al., 2001; Auer and Benton, 2016; Baker et al., 2001; Billeter et al., 2006; Clyne and Miesenböck, 2008; Dauwalder, 2011; Demir and Dickson, 2005; Evans and Cline, 2007; Goodwin et al., 2000; Hall, 1994; Inoue et al., 1992; Ito et al., 1996; Kimura et al., 2019; Kvitsiani and Dickson, 2006; Neville et al., 2014; Nojima et al., 2014; Pavlou et al., 2016; von Philipsborn et al., 2014; Rezával et al., 2014, 2016; Rideout et al., 2007, 2010; Ryner et al., 1996; Sato et al., 2019; Shirangi et al., 2016; Wang et al., 2020), less is known about sex differences in neurons that regulate physiology and metabolism. Indeed, while many studies have identified neurons that regulate fat metabolism (Al-Anzi and Zinn, 2018; Al-Anzi et al., 2009; Chung et al., 2017; Li et al., 2016; May et al., 2020; Min et al., 2016; Mosher et al., 2015; Zhan et al., 2016), these studies were conducted in single- or mixed-sex populations. Because male-female differences in neuron number (Billeter et al., 2006; Castellanos et al., 2013; Demir and Dickson, 2005; Garner et al., 2018; Lee and Hall, 2001; Rideout et al., 2007, 2010; Robinett et al., 2010; Taylor and Truman, 1992), morphology (Cachero et al., 2010; Kimura et al., 2019), activity (Guo et al., 2016), and connectivity (Cachero et al., 2010; Nojima et al., 2021) have all been described across the brain and ventral nerve cord (Mellert et al., 2010, 2016), a detailed analysis of neuronal populations that influence metabolism will be needed in both sexes to understand how neurons contribute to the sex-specific regulation of metabolism and physiology. Indeed, while our identification of a role for APC sexual identity in regulating the male-female difference in fat storage represents a significant step forward in understanding how sex differences in neurons influence metabolic traits, more knowledge is needed of sexual dimorphism in this critical neuronal subset.

An obvious starting point for learning more about the sex-specific regulation of fat storage by the APCs is to examine how sexual identity influences known APC regulatory mechanisms. For example, there are physical connections between corazonin- and neuropeptide F (NPF; FBgn0027109)-positive (CN) neurons and APCs in adult male flies (Oh et al., 2019), and between the APCs and a bursicon-*α*-responsive subset of DLgr2 neurons in females (Scopelliti et al., 2019). These connections inhibit APC activity: CN neurons inhibit APC activity in response to high hemolymph sugar levels (Oh et al., 2019), whereas binding of bursicon-*α* to DLgr2 neurons inhibits APC activity in nutrient-rich conditions (Scopelliti et al., 2019). Future studies will therefore need to determine whether these physical connections exist in both sexes. Male-female differences in circulating factors that regulate the APCs may also exist. For example, gut-derived Allatostatin C (AstC; FBgn0032336) binds its receptor on the APCs to trigger Akh release; however, loss of AstC affects fat metabolism and starvation resistance in females but not in males (Kubrak et al., 2020). This suggests that sex differences in AstC production or release may exist. Additionally, gut-derived NPF binds to its receptor on APCs to inhibit Akh release (Yoshinari et al., 2021). Another circulating factor that may influence the sex difference in fat storage is skeletal muscle-derived unpaired 2 (upd2; FBgn0030904), which regulates hemolymph Akh levels to maintain diurnal fat metabolism (Zhao and Karpac, 2017). Given that circulating peptides such as Allatostatin A (AstA; FBgn0015591), *Drosophila* insulin-like peptides (Dilps), and activin ligands also influence Akh pathway activity (Ahmad et al., 2020; Hentze et al., 2015; Post et al., 2019; Song et al., 2017), a systematic survey of circulating factors that modulate Akh production, release, and Akh pathway activity in each sex will be needed to fully understand the sex-specific regulation of fat storage.

In addition to fat metabolism, it will be important to extend our understanding of how sex-specific Akh regulation affects additional Akh-regulated phenotypes. For example, Akh has been linked with the regulation of lifespan (Bednářová et al., 2018; Liao et al., 2021), starvation resistance (Isabel et al., 2005; Kubrak et al., 2020; Mochanová et al., 2018), locomotion (Isabel et al., 2005; Lee and Park, 2004), immune responses (Adamo et al., 2008), cardiac function (Isabel et al., 2005; Noyes et al., 1995), oxidative stress responses (Gáliková et al., 2015), and fertility (Liao et al., 2021). Yet, most studies were performed in mixed- or single-sex populations. This suggests additional work is needed to determine how changes to Akh pathway function affect physiology, development, and life history in both sexes. Importantly, the lessons we learn may also extend to other species. Akh signalling is highly conserved across invertebrates (Gäde and Auerswald, 2003; Lorenz and Gäde, 2009; Staubli et al., 2002b), and is functionally similar to the mammalian *β*-adrenergic and glucagon systems (Grönke et al., 2007; Lee and Park, 2004; Staubli et al., 2002b). Because sex-specific regulation of both glucagon and the *β*-adrenergic systems have been described in mammalian models and in humans (Al-Gburi et al., 2017; Bell et al., 2001; Bilginoglu et al., 2007; Brooks et al., 2015; Claustre et al., 1980; Dart et al., 2002; Davis et al., 2000; Drake et al., 1998; Freedman et al., 1987; Hinojosa-Laborde et al., 1999; Hoeker et al., 2014; Hogarth et al., 2007; Lafontan et al., 1997; Luzier et al., 1998; McIntosh et al., 2011; Ng et al., 1993), detailed studies on sex-specific Akh regulation and function in flies may provide vital clues into the mechanisms underlying male-female differences in physiology and metabolism in other animals.

## MATERIALS AND METHODS

### Data availability

Details of all statistical tests and *p-*values are in Supplementary file 1. All raw data generated in this study are in Supplementary file 2. All primer sequences are in Supplementary file 3. Fly food media recipe is in Supplementary file 4. Original image files for all images in this study are in their respective Source Data files.

### Fly husbandry

Fly stocks were maintained at 25°C in a 12:12 light:dark cycle. All larvae were reared at a density of 50 larvae per 10 ml of fly media (recipe in Supplementary file 4). Males and females were separated either as early pupae by gonad size, or late pupae by the presence of sex combs. Sex-transformed males and females were distinguished by the presence (males) or absence (females) of B^S^Y. Single-sex groups of 20 pupae were transferred to damp filter paper within a food vial until eclosion. Unless otherwise stated, all experiments used 5- to 7-day-old flies.

### Fly strains

We obtained the following strains from the Bloomington *Drosophila* Stock Center: *Canton-S* (#64349), *w^1118^* (#3605), *UAS-nGFP* (#4775), *UAS-Akh-RNAi* (#27031), *UAS-tra^F^* (#4590), *tra^1^* (#675), *Df(3L)st-j7* (#5416), *UAS-NaChBac* (#9468), *UAS-Kir2.1* (#6595), *UAS-reaper* (#5823), *UAS-CaLexA* (#66542). We obtained *Akh^A^*, *AkhR^rev^*, *AkhR^1^*, *bmm^1^*, and *AkhR^1^;bmm^1^* as kind gifts from Dr. Ronald Kühnlein (Gáliková et al., 2015; Grönke et al., 2005, 2007), *tra^KO^* and *tra^F K-IN^* as kind gifts from Dr. Irene Miguel-Aliaga (Hudry et al., 2016, 2019), and *Mex-GAL4* as a kind gift from Dr. Claire Thomas (Phillips and Thomas, 2006). The following GAL4 lines were used for tissue-specific expression: *da-GAL4* (ubiquitous), *cg-GAL4* (fat body), *r4-GAL4* (fat body), *Lsp2-GAL4* (fat body), *Myo1A-GAL4* (enterocytes), *Mex-GAL4* (enterocytes), *dMef2-GAL4* (skeletal muscle), *repo-GAL4* (glia), *elav-GAL4* (neurons), *c587-GAL4* (somatic cells of the gonad), *tj-GAL4* (somatic cells of the gonad), *nos-GAL4* (germ cells of the gonad), *dimmed-GAL4* (peptidergic neurons), *TH-GAL4* (dopaminergic neurons), *Tdc2-GAL4* (octopaminergic neurons), *VT030559-GAL4* (mushroom body neurons), *dilp2-GAL4* (insulin-producing cells), *Akh-GAL4* (Akh-producing cells). All transgenic stocks were backcrossed into a *w^1118^* background for a minimum of 5 generations.

### Adult weight

To measure adult weight, groups of ten flies were weighed in 1.5 ml microcentrifuge tubes on an analytical balance (Mettler-Toledo, ME104).

### RNA extraction, cDNA synthesis, and qPCR

One biological replicate consisted of five flies homogenized in 200 μl of Trizol. RNA was extracted following manufacturer’s instructions, as previously described (Wat et al., 2020). cDNA was synthesized from RNA using the Quantitect Reverse Transcription Kit (Qiagen, 205311). qPCR was used to quantify relative mRNA transcript levels as previously described (Wat et al., 2020).

See Supplementary file 3 for a full list of primers.

### Whole-body triglyceride measurements

One biological replicate consisted of five flies homogenized in 200 μl of 0.1% Tween (AMresco, 0777-1L) in 1X PBS using 50 μl of glass beads (Sigma, 11079110) agitated at 8 m/s for 5 seconds (OMNI International Bead Ruptor 24). Assay was performed according to established protocols (Tennessen et al., 2014) as previously described (Wat et al., 2020).

### Western blotting

One biological replicate consisted of ten flies homogenized in extraction buffer (females=200 μl, males=125 μl) containing 20 mM Hepes (pH 7.8), 450 mM NaCl, 25% glycerol, 50 mM NaF, 0.2 mM EDTA, 0.5% Triton X-100, 1 mM PMSF, 1 mM DTT, 1X cOmplete Protease Inhibitor Cocktail (Roche), and 1X PhosSTOP (Roche) using 50 μl of glass beads (Sigma, 11079110) agitated at 8 m/s for 5 seconds (OMNI International Bead Ruptor 24). Samples were incubated on ice for 5 min before cellular debris was pelleted by centrifugation at 10,000 rpm for 5 min at 4°C and supernatant was removed (Thermo Scientific, Heraeus Pico 21 centrifuge). Centrifugation was repeated twice more to remove fat from the samples. Protein concentration of each sample was determined by a Bradford Assay (Bio-Rad, 550-0205); 20 μg of protein per sample was loaded onto a 12% SDS-PAGE gel. Immunoblotting was performed as previously described (Millington et al., 2021). Primary antibodies used were rabbit anti-p-Ire1 (1:1000; Abcam #48187) and mouse anti-actin (1:100; Santa Cruz #sc-8432).

Secondary antibodies used were goat anti-rabbit (1:5000; Invitrogen #65-6120) and horse anti-mouse (1:2000; Cell Signaling #7076).

### APC measurements

To isolate the APCs, individual flies were anesthetized on ice, and the brain and foregut were removed in cold 1X PBS. Samples were fixed in 4% paraformaldehyde for 30 min, followed by two 30 min washes in cold 1X PBS. Samples were incubated with Hoechst (Sigma, 33342) at a concentration of 1:500 for 5 min and mounted in SlowFade Diamond Antifade Mountant (ThermoFisher, S36967). Images were captured using a Leica TCS SP5 Confocal Microscope and processed using Fiji (ImageJ; Schindelin et al., 2012). To visualize APC neuronal activity (*Akh-GAL4>UAS-CaLexA*), the mean GFP intensity of one APC cluster was quantified by measuring average pixel intensity within the region of interest using Fiji (ImageJ; Schindelin et al., 2012). To determine APC cell number (*Akh-GAL4>UAS-nGFP*), GFP punctae were counted manually using Fiji (ImageJ; Schindelin et al., 2012). One biological replicate consists of one cluster of APCs, where only one APC cluster was measured per individual.

### Capillary Feeder Assay

One biological replicate consisted of ten flies placed into a specialized 15 ml conical vial with access to 2 capillary tubes. Capillary tubes were filled with fly food media containing 5% sucrose, 5% yeast extract, 0.3% propionic acid, and 0.15% nipagin. Approximately 0.5 μl of mineral oil was added to the top of each capillary tube to prevent evaporation. All vials were placed into fitted holes in the lid of a large plastic container. A shallow layer of water was poured into the base of the container to maintain high humidity throughout the experiment. The meniscus of the fly food media was marked before the start of the experiment and again after 24 hr. The distance between the marks is used to quantify the volume of fly food media that was consumed (1 mm=0.15 μl). The volume of fly food consumed was normalized to the weight of individual flies (protocol adapted from Stafford et al., 2012).

### Male fertility

One singly-housed male was placed with a group of three virgin *Canton-S (CS*) females and allowed to interact for 60 min. At 10 min intervals during the 60 min observation period, we recorded whether a copulating male-female pair was present in the vial. After the 60 min observation period, the male was removed from the vial and the females were allowed to lay eggs for 72 hr (flies were transferred to new food every 24 hr). After 72 hr, the females were removed and progeny were allowed to develop.

After 10 days, we counted the number of pupae in each vial. For the 24 hr mating assay, one singly-housed male was allowed to interact with three virgin *CS* females for 24 hr before the male was removed and females were allowed to lay eggs for 72 hr as described above.

### Female fecundity

One virgin female was placed with a group of three virgin *CS* males for 24 hr. The females were then transferred onto fresh food every 24 hr for 3 days and the number of pupae were counted as described above.

### Starvation resistance

5-day-old flies were transferred to vials containing 2 ml of starvation media (0.7% agar in 1X PBS). The number of deaths was recorded every 12 hr until no living flies remained in the vial.

### Lifespan

Flies were transferred to new vials with 2 ml of fresh food every 2-3 days until no living flies remained in the vial. Deaths were recorded when the flies were transferred.

### Statistics and data presentation

All figures and data were generated and analyzed using GraphPad Prism (v9.1.2). For experiments with 2 groups, a Student’s *t-*test was performed. For experiments with 3 or more groups, a one-way ANOVA with Tukey HSD post-hoc test was performed. For fat breakdown experiments, a two-way ANOVA was used to determine the interaction between genotype and time. Starvation resistance and lifespan statistics were performed using RStudio and a script for a Log-rank test with Bonferroni’s correction for multiple comparisons. Note, the lowest *p*-value given by RStudio is 2.0 x 10^-16^. The below packages and script were used: library (“survminer”) library (“survival”) data <- read.csv(“xxx.csv”) survfit(Surv(time, event) ∼ genotype, data) pairwise_survdiff(Surv(time, event) ∼ genotype, data, p.adjust.method = “bonferroni”) summary (data)

## ACKNOWLEDGEMENTS

We would like to thank Dr. Ronald Kühnlein for *Akh^A^, AkhR^rev^, AkhR^1^, bmm^1^, and AkhR^1^;bmm^1^* fly strains. We would also like to thank Dr. Irene Miguel-Aliaga for the *tra^KO^* and *tra^F K-IN^* strains, and Dr. Claire Thomas for *Mex-GAL4.* Stocks obtained from the Bloomington *Drosophila* Stock Center (NIH P40OD018537) were used in this study. We thank the TRiP at Harvard Medical School (NIH/NIGMS R01-GM084947) for providing transgenic RNAi fly stocks and/or plasmid vectors used in this study. We acknowledge critical resources and information provided by FlyBase (Thurmond et al., 2019); FlyBase is supported by a grant from the National Human Genome Research Institute at the U.S. National Institutes of Health (<u>U41 HG000739</u>) and by the British Medical Research Council (<u>MR/N030117/1</u>). Funding for this study was provided by grants to EJR from the Canadian Institutes for Health Research (CIHR, PJT-153072), CIHR Sex and Gender Science Chair Program (GS4-171365), Natural Sciences and Engineering Research Council of Canada (NSERC, RGPIN-2016-04249), Michael Smith Foundation for Health Research (16876), and the Canadian Foundation for Innovation (JELF-34879). LWW was supported by a British Columbia Graduate Scholarship Award and a 1-year CELL Fellowship from UBC. JWM and PB were each supported by a 4-year CELL Fellowship from UBC. We acknowledge that our research takes place on the traditional, ancestral, and unceded territory of the Musqueam people; a privilege for which we are grateful.

## SUPPLEMENTAL FIGURE LEGENDS

**Figure 1 - figure supplement 1.**
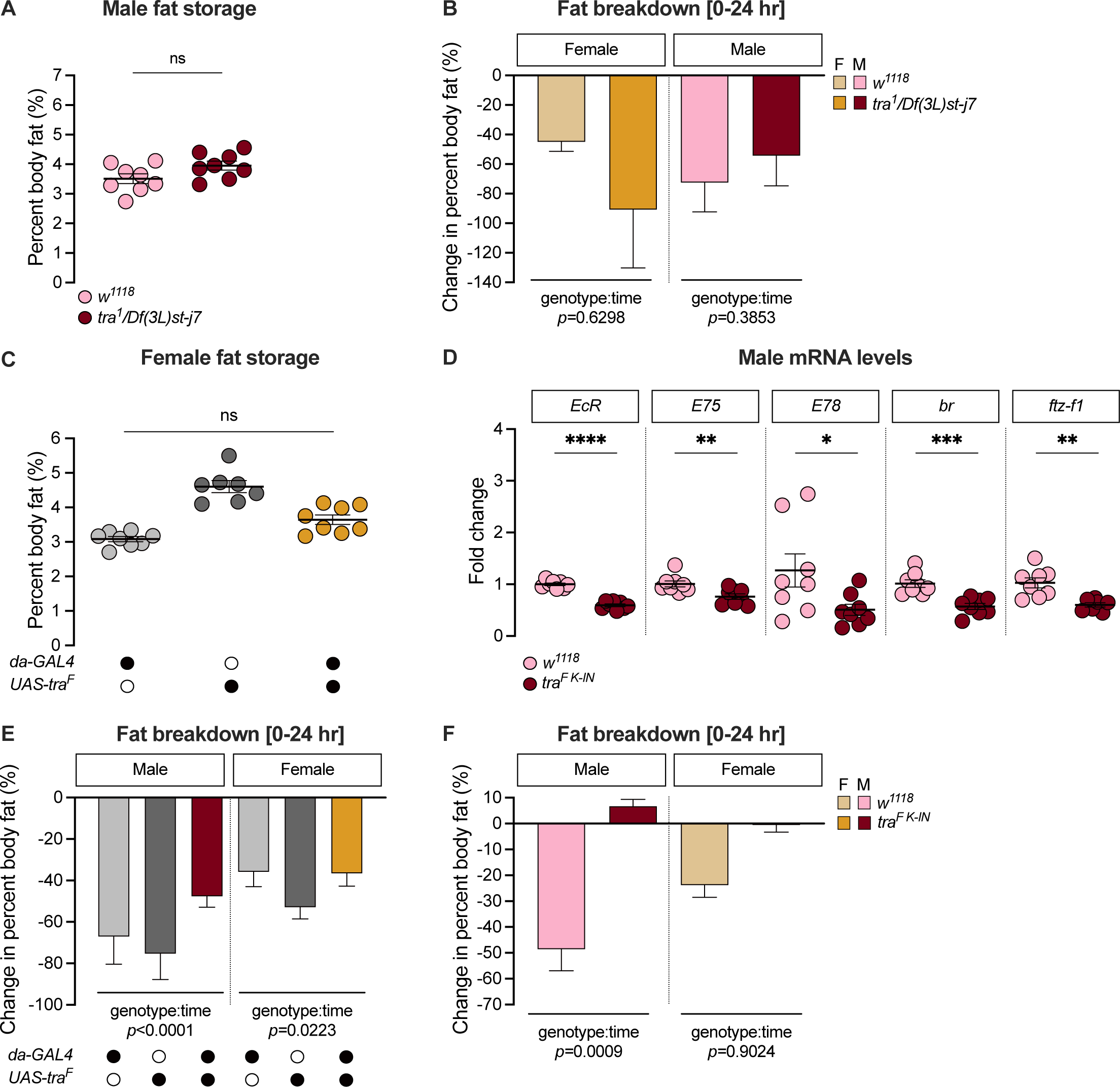
Elucidating *transformer*’s effect on sex differences in fat metabolism. (A) Whole-body triglyceride levels were not significantly different between *tra^1^/Df(3L)st-j7* males and *w^1118^* controls (*p*=0.0685; Student’s *t*-test). *n*=8 biological replicates. (B) The reduction in whole-body triglyceride levels post-starvation was not significantly different between *tra^1^/Df(3L)st-j7* animals and sex-matched *w^1118^* controls between 0-24 hr post-starvation (genotype:time *p*=0.6298 [female], *p*=0.3853 [male]; two-way ANOVA per sex). *n*=7-8 biological replicates. (C) Whole-body triglyceride levels in *da-GAL4>UAS-tra^F^ fe*males were intermediate between *da- GAL4>+* and *+>UAS-tra^F^* control females, indicating no overall effect of Tra overexpression in females (*p*=0.0160 and *p*=0.0002 respectively; one-way ANOVA followed by Tukey’s HSD). *n*=7-8 biological replicates. (D) Whole-body mRNA levels of ecdysone responsive genes were not higher in *tra^F K-IN^* males compared to *w^1118^* control males (*Ecdysone receptor* (*EcR*)*: p*<0.0001; *Ecdysone-induced protein 75B (E75): p*=0.0072; *Ecdysone-induced protein 78C (E78): p*=0.0408; *broad (br): p*=0.0003; *ftz transcription factor 1* (*ftz-f1*)*: p*=0.002; Student’s *t*-test for each gene). *n*=7-8 biological replicates. (E) The reduction in whole-body triglyceride levels between 0-24 hr post-starvation was significantly smaller in *da-GAL4>UAS-tra^F^* males compared with *da- GAL4>+* and *+>UAS-tra^F^* control males (genotype:time *p*<0.0001; two-way ANOVA). The post-starvation reduction in triglyceride levels in *da-GAL4>UAS-tra^F^* females was intermediate between both *da-GAL4>+* and *+>UAS-tra^F^* controls, suggesting no overall effect of Tra overexpression on female fat storage (genotype:time *p*=0.0223; two-way ANOVA per sex). *n*=7-8 biological replicates. (F) The reduction in whole-body triglyceride levels between 0-24 hr post-starvation was significantly lower in *tra^F K-IN^* males, but not females, compared with sex-matched *w^1118^* controls (genotype:time *p*=0.0009 [male], *p*=0.9024 [female]; two-way ANOVA per sex). *n*=7-8 biological replicates. F indicates female, M indicates male. Black circles indicate the presence of a transgene and open circles indicate the lack of a transgene. * indicates *p*<0.05, ** indicates *p*<0.01, *** indicates *p*<0.001, **** indicates *p*<0.0001, ns indicates not significant; error bars represent SEM except for graphs displaying fat breakdown where error bars represent COE.

**Figure 2 - figure supplement 1.**
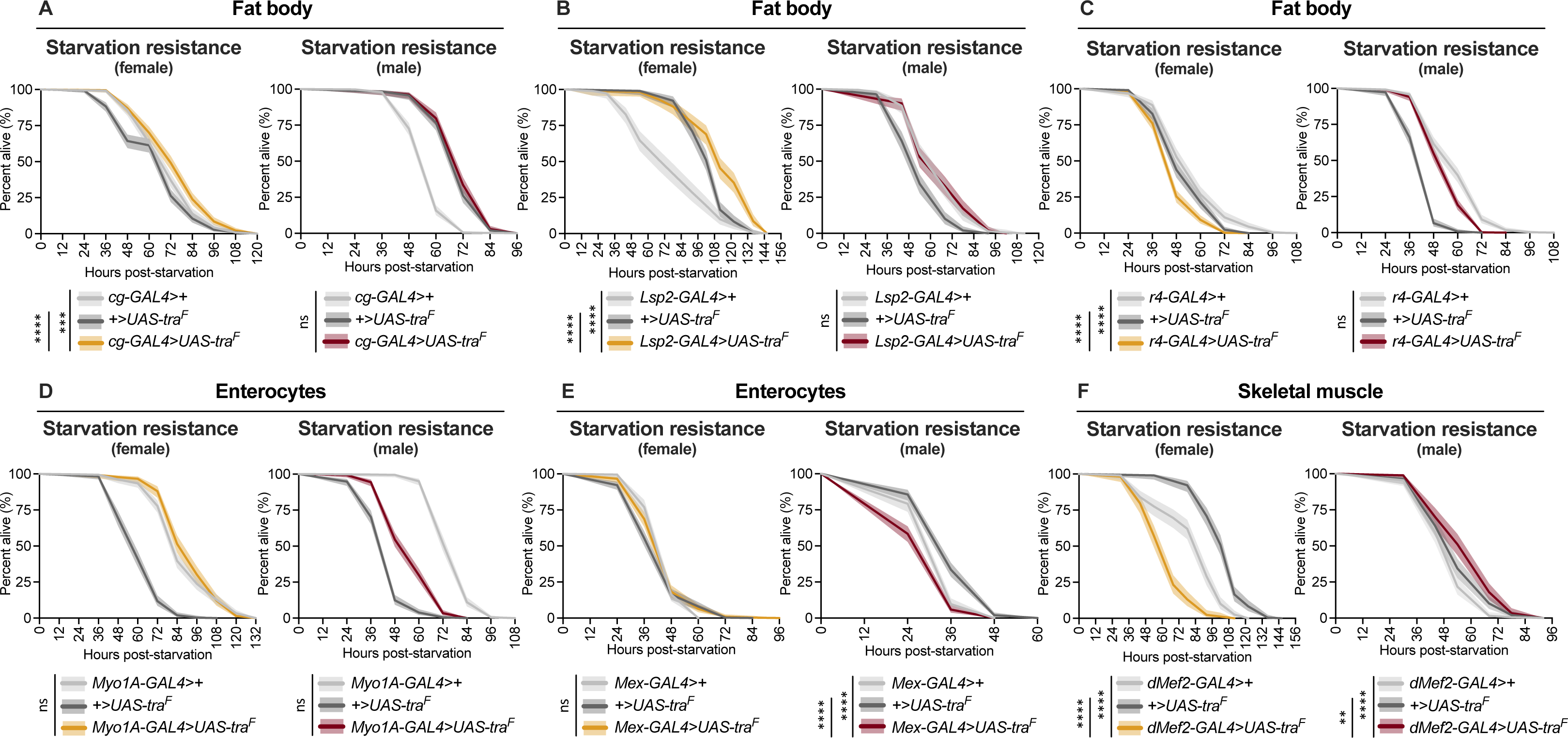
Effect of *transformer* gain in multiple cell types and tissues on starvation resistance. (A-F) Limited to no effects of Tra overexpression on starvation resistance were observed for female fat body, muscle, and gut (for individual *p*-values see Supplementary file 1; log-rank test with Bonferroni’s correction for multiple comparisons). In males, Tra overexpression in the fat body and gut caused no extension of starvation resistance, with only a minor extension observed upon Tra overexpression in muscle (for individual *p*-values see Supplementary file 1; log-rank test with Bonferroni’s correction for multiple comparisons). (A) *n*=397-413 females, *n*=295-431 males. (B) *n*=187-206 females, *n*=198-202 males. (C) *n*=293-402 females, *n*=330-452 males. (D) *n*=268-383 females, *n*=363-409 males. (E) *n*=226-374 females, *n*=250-362 males. (F) *n*=168-206 females, *n*=178-198 males. ** indicates *p*<0.01, *** indicates *p*<0.001, **** indicates *p<*0.0001, ns indicates not significant; shaded areas represent the 95% confidence interval.

**Figure 2 - figure supplement 2.**
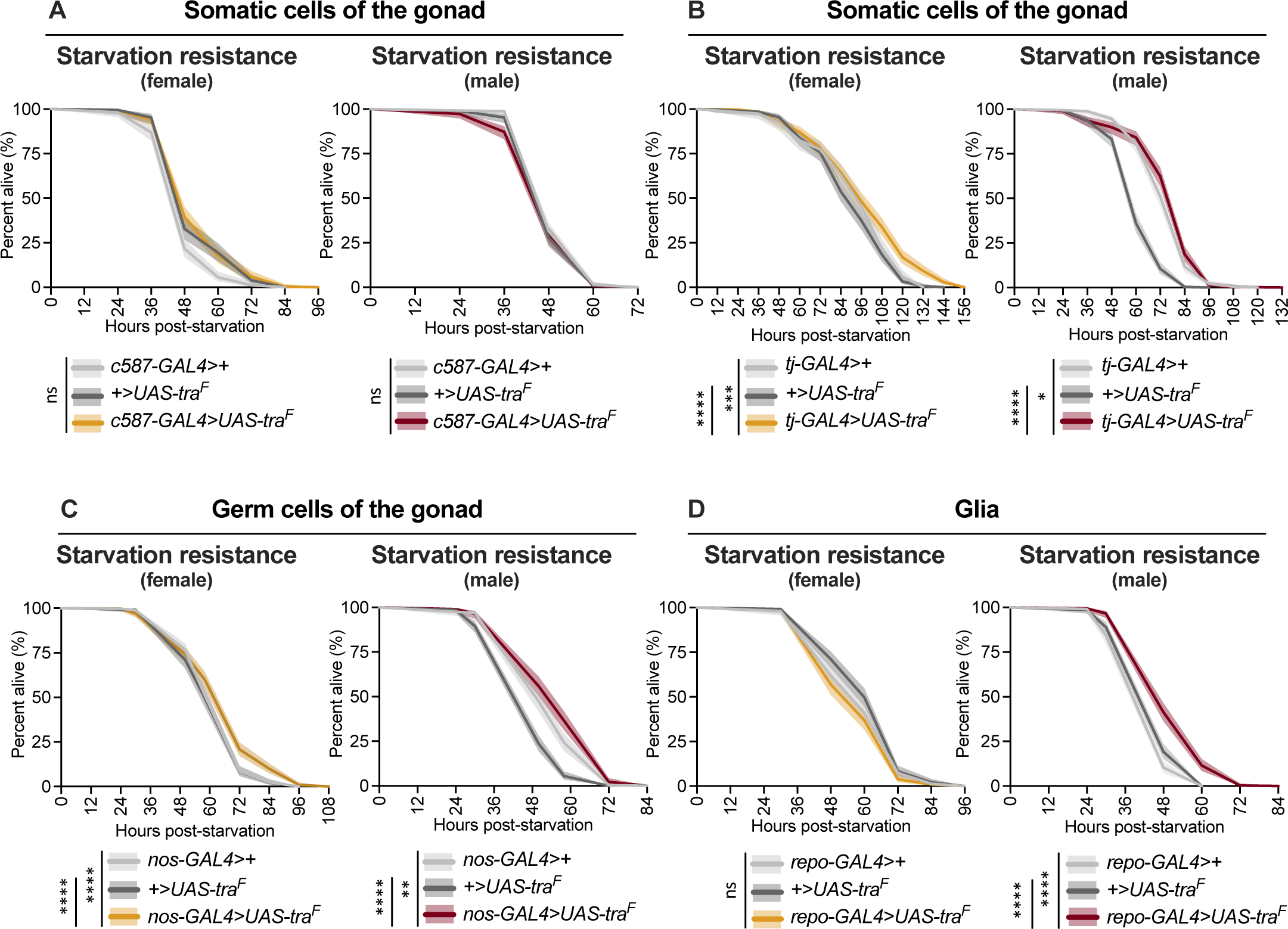
Effect of *transformer* gain in additional cell types and tissues on starvation resistance. (A-D) Limited to no effects of Tra overexpression were observed upon Tra expression in the female gonad and glia (for individual *p*-values see Supplementary file 1; log-rank test with Bonferroni’s correction for multiple comparisons). Limited to no effects of Tra overexpression were observed upon Tra expression in the male gonad and glia (for individual *p*-values see Supplementary file 1; log-rank test with Bonferroni’s correction for multiple comparisons). (A) *n*=232-268 females, *n*=318-400 males. (B) *n*=234-349 females, *n*=327-349 males. (C) *n*=374-442 females, *n*=293-364 males. (D) *n*=300-374 females, *n*=329-355 males. * indicates *p*<0.05, ** indicates *p*<0.01, *** indicates *p*<0.001, **** indicates *p*<0.0001, ns indicates not significant; shaded areas represent the 95% confidence interval.

**Figure 2 - figure supplement 3.**
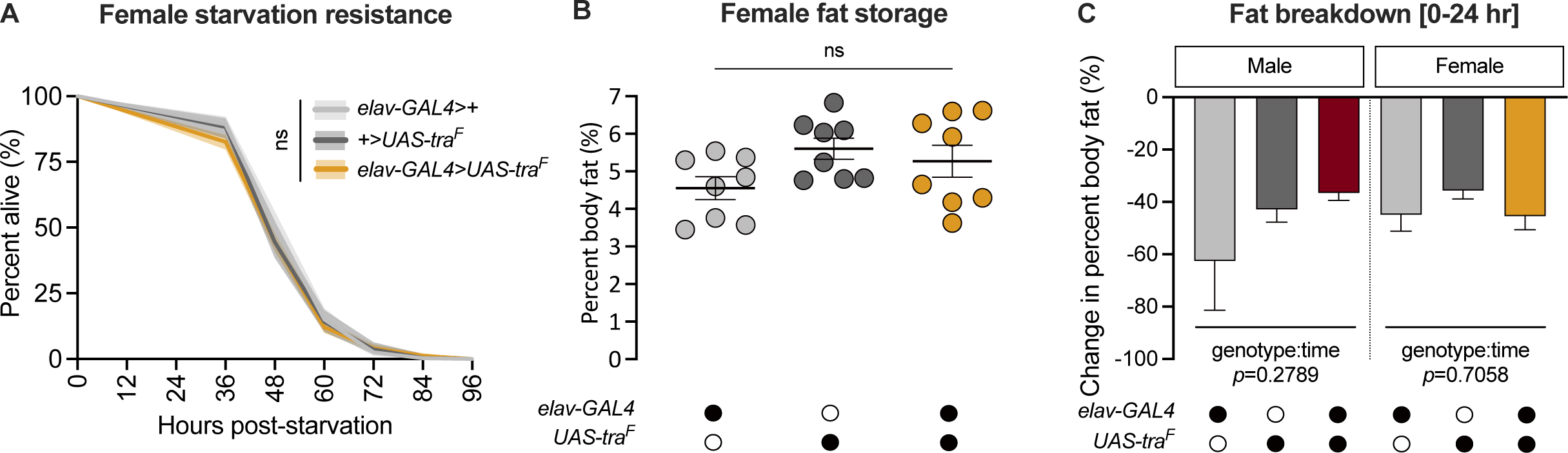
Gain of *transformer* function in neurons does not affect fat breakdown. (A) Starvation resistance in *elav-GAL4>UAS-tra^F^* females was not significantly different compared with *elav-GAL4>+* and *+>UAS-tra^F^* controls (*p*=0.3 and *p*=1, respectively; log-rank test with Bonferroni’s correction for multiple comparisons). *n*=318-749 females. (B) Whole-body triglyceride levels were not significantly different between *elav-GAL4>UAS-tra^F^* females and *elav-GAL4>+* and *+>UAS-tra^F^* controls (*p*=0.3224 and *p*=0.7754 respectively; one-way ANOVA followed by Tukey’s HSD). *n*=8 biological replicates. (C) The reduction in whole-body triglyceride levels post-starvation was not significantly different between *elav-GAL4>UAS-tra^F^* flies and sex-matched *elav-GAL4>+* and *+>UAS-tra^F^* controls (genotype:time *p*=0.2789 [male], *p*=0.7058 [female]; two-way ANOVA per sex). *n*=7-8 biological replicates. Black circles indicate the presence of a transgene and open circles indicate the lack of a transgene; ns indicates not significant; shaded areas represent the 95% confidence interval; error bars represent SEM except for graphs displaying fat breakdown where error bars represent COE.

**Figure 2 - figure supplement 4.**
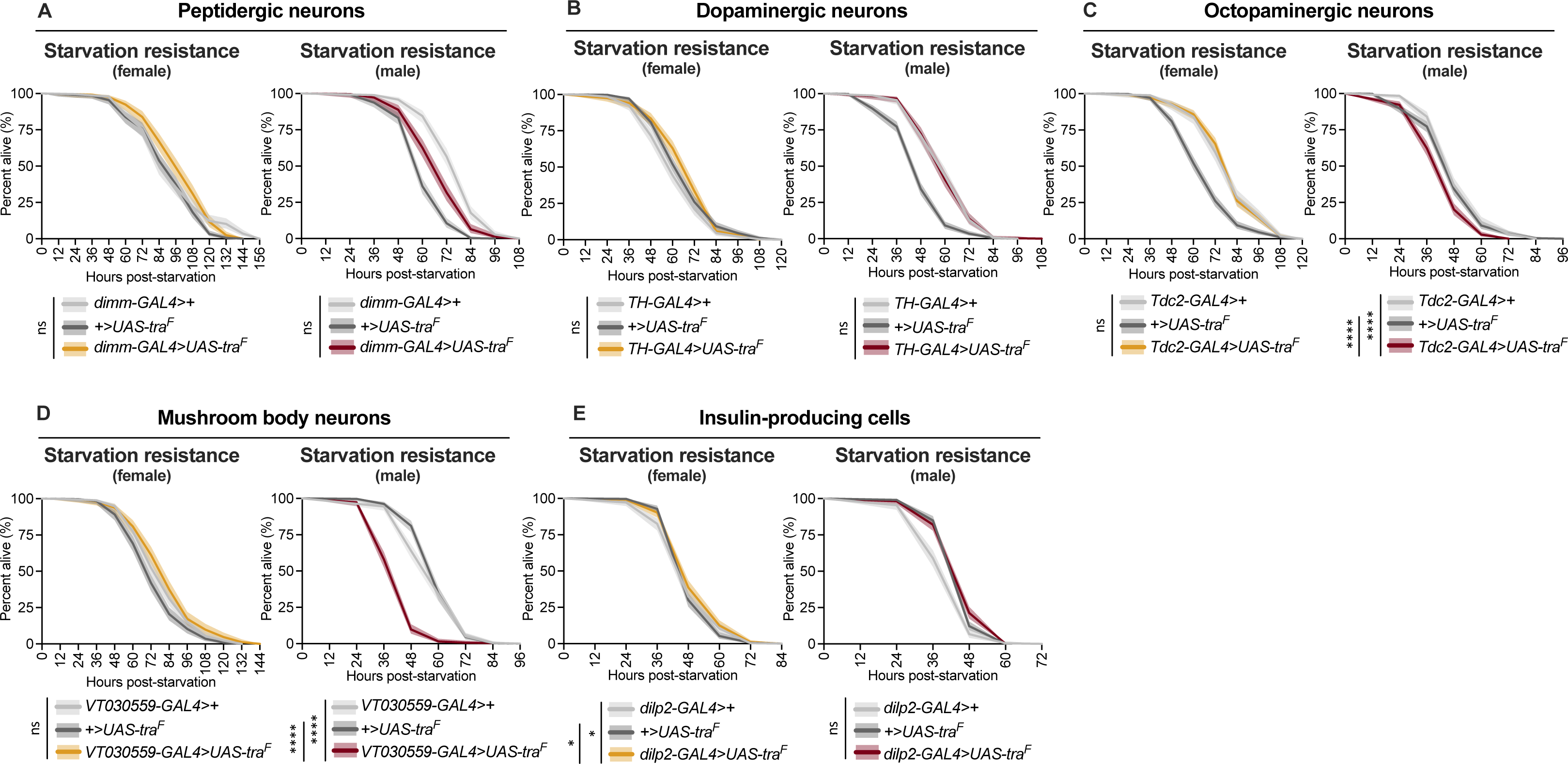
Effect of *transformer* gain in multiple neuronal subsets on starvation resistance. (A-E) Limited to no effects of Tra overexpression in several subsets of neurons on starvation resistance in females (for individual *p*-values see Supplementary file 1; log-rank test with Bonferroni’s correction for multiple comparisons). In males, Tra overexpression in several subsets of neurons caused no extension of starvation resistance (for individual *p*-values see Supplementary file 1; log-rank test with Bonferroni’s correction for multiple comparisons). (A) *n*=248-333 females, *n*=249-333 males. (B) *n*=322-484 females, *n*=314-516 males. (C) *n*=282-478 females, *n*=364-516 males. (D) *n*=256-343 females, *n*=326-392 males. (E) *n*=326-390 females, *n*=285-466 males. * indicates *p*<0.05, **** indicates *p*<0.0001, ns indicates not significant; shaded areas represent the 95% confidence interval.

**Figure 2 - figure supplement 5.**
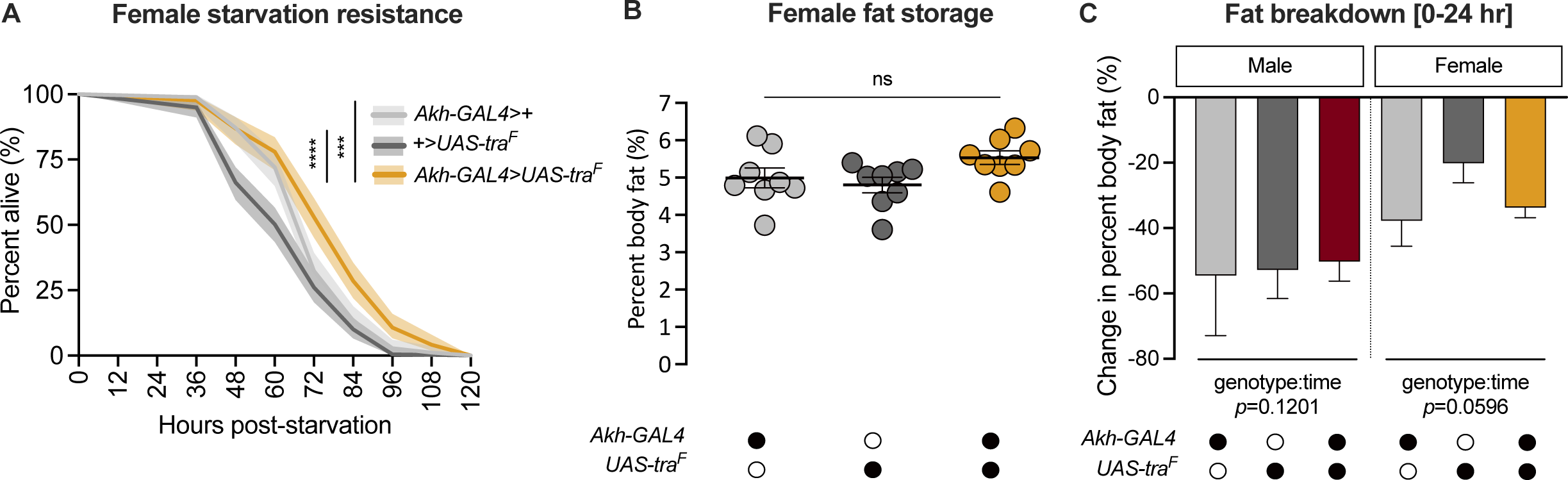
Gain of *transformer* function in Akh-producing cells does not affect fat breakdown. (A) Starvation resistance was significantly extended in *Akh-GAL4>UAS-tra^F^* females compared with *Akh-GAL4>+* and *+>UAS-tra^F^* controls (*p=*0.00033 and *p*=9.4x10^-11^ respectively; log-rank test with Bonferroni’s correction for multiple comparisons). *n*=168-219 females. (B) Whole-body triglyceride levels were not significantly different between *Akh-GAL4>UAS-tra^F^* females and *Akh-GAL4>+* and *+>UAS-tra^F^* controls (*p*=0.2195 and *p*=0.0731 respectively; one-way ANOVA followed by Tukey’s HSD). *n*=8 biological replicates. (C) The reduction in whole-body triglyceride levels post-starvation was not significantly different between *Akh-GAL4>UAS-tra^F^* animals and sex-matched *Akh-GAL4>+* and *+>UAS-tra^F^* controls (genotype:time *p*=0.1201 [males], *p*=0.0596 [female]; two-way ANOVA per sex). *n*=8 biological replicates. Black circles indicate the presence of a transgene and open circles indicate the lack of a transgene; *** indicates *p*<0.001, **** indicates *p*<0.0001, ns indicates not significant; shaded areas represent the 95% confidence interval; error bars represent SEM except for graphs displaying fat breakdown where error bars represent COE.

**Figure 3 - figure supplement 1.**
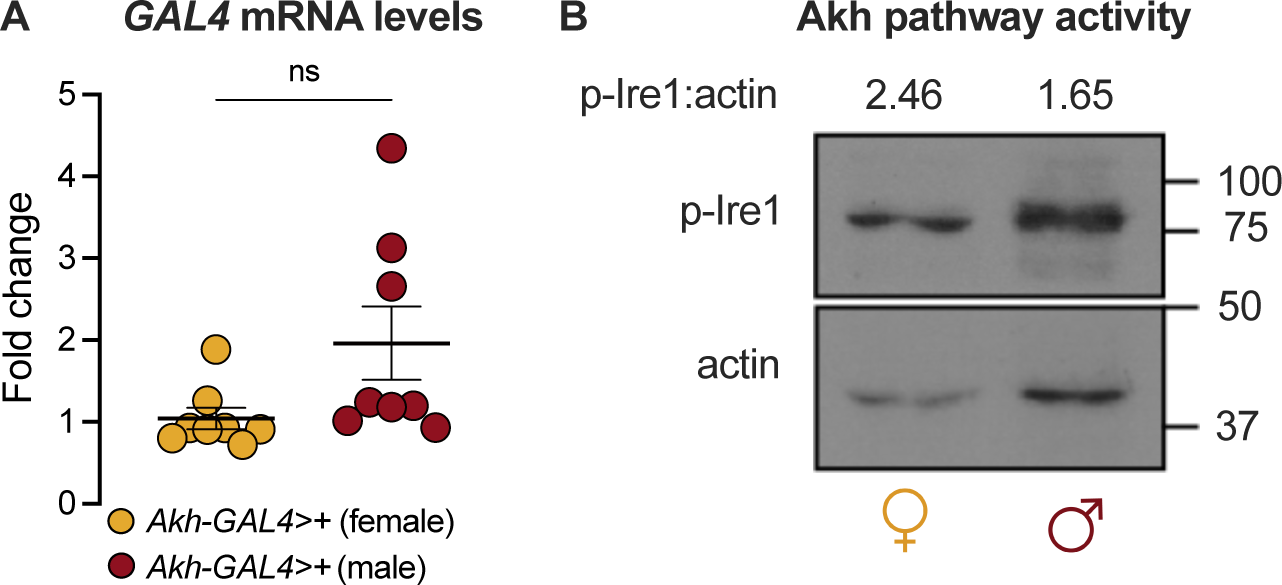
*Akh-GAL4* drives equivalent *GAL4* mRNA levels in both sexes. (A) Whole-body *GAL4* mRNA levels were not significantly different between *Akh-GAL4>+* females and males (*p*=0.0687; Student’s *t*-test). *n*=8 biological replicates. (B) Whole-body p-Ire1 levels were not higher in *w^1118^* males compared with *w^1118^* females in one biological replicate. ns indicates not significant; error bars represent SEM. Original image for Figure 3 – figure supplement 1B is found in Figure 3 – figure supplement 1 - Source Data 1.

**Figure 4 - figure supplement 1.**
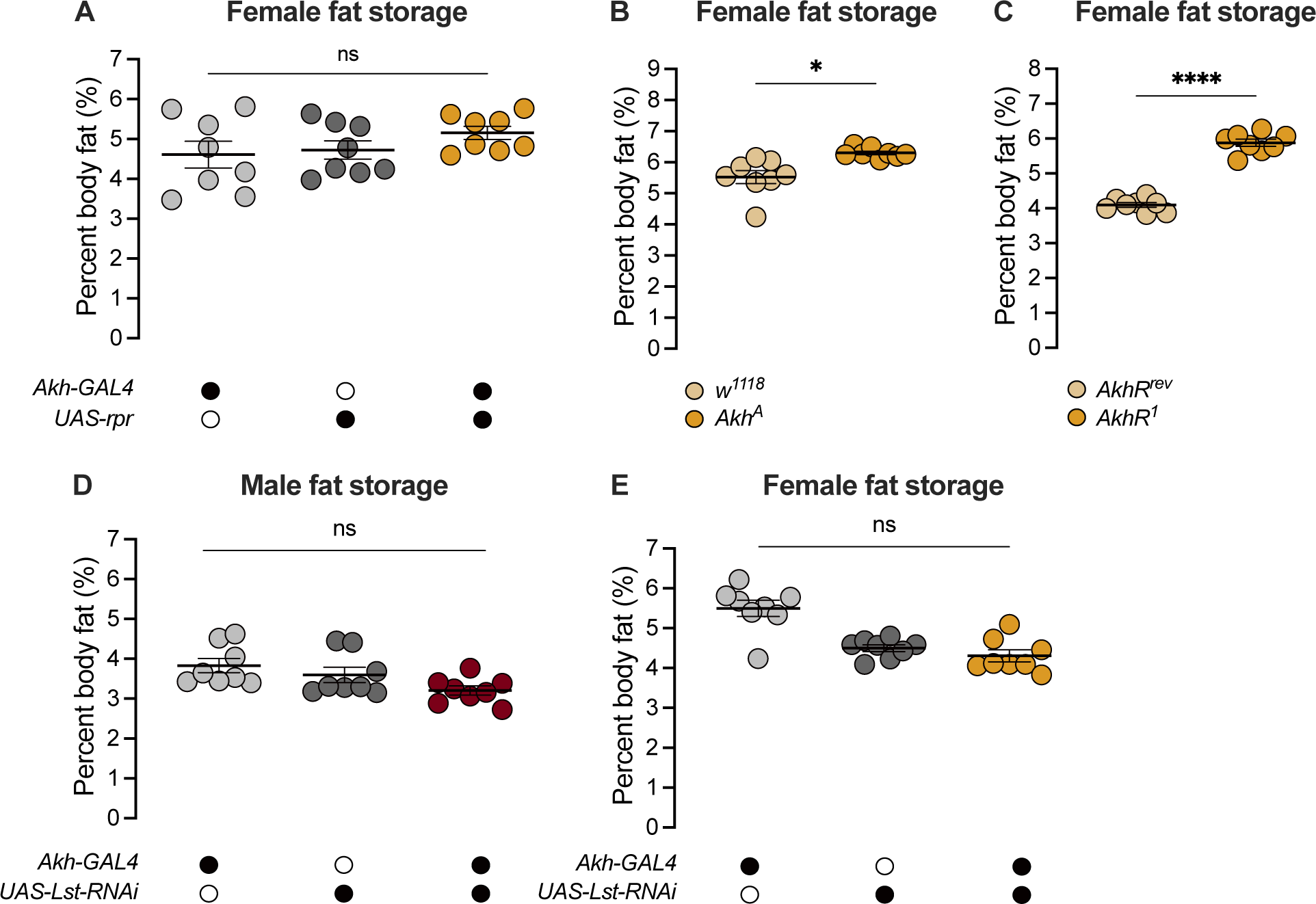
APC-derived Limostatin does not regulate the sex difference in fat storage. (A) Whole-body triglyceride levels were not significantly different between *Akh-GAL4>UAS-reaper (rpr)* females and *Akh-GAL4>+* and *+>UAS- rpr* controls (*p*=0.3024 and *p*=0.4673 respectively; one-way ANOVA followed by Tukey’s HSD). *n*=8 biological replicates. (B) Whole-body triglyceride levels were significantly higher in *Akh^A^* females compared with *w^1118^* controls (*p*=0.0152; one-way ANOVA followed by Tukey’s HSD). *n*=8 biological replicates. (C) Whole-body triglyceride levels were significantly higher in *AkhR^1^* females compared with *AkhR^rev^* control females (*p*<0.0001; one-way ANOVA followed by Tukey’s HSD). *n*=8 biological replicates. (D) Whole-body triglyceride levels in *Akh-GAL4>UAS-Limostatin (Lst)-RNAi* males were not significantly different from both *Akh-GAL4>+* and *+>UAS-Lst-RNAi* controls (*p*=0.0357 and *p*=0.2364 respectively; one-way ANOVA followed by Tukey’s HSD). *n*=8 biological replicates. (E) Whole-body triglyceride levels in *Akh-GAL4>UAS-Lst-RNAi* females were not significantly different from both *Akh-GAL4>+* and *+>UAS-Lst-RNAi* controls (*p*<0.0001 and *p*=0.6656 respectively; one-way ANOVA followed by Tukey’s HSD). *n*=8 biological replicates. Black circles indicate the presence of a transgene and open circles indicate the lack of a transgene. * indicates *p*<0.05, **** indicates *p*<0.0001, ns indicates not significant; error bars represent SEM.

**Figure 4 - figure supplement 2.**
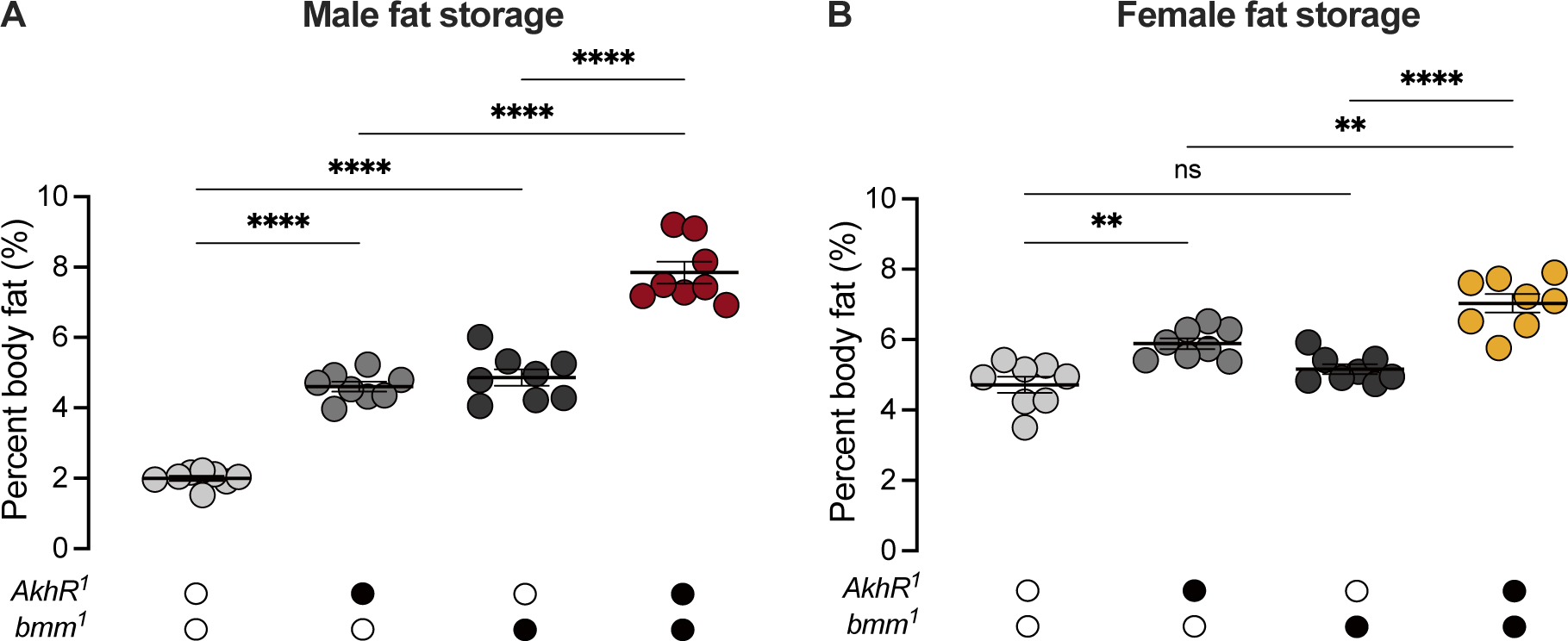
Akh and *brummer* operate in parallel pathways to regulate the sex difference in fat storage. (A) Whole-body triglyceride levels were significantly higher in *AkhR^1^* and *bmm^1^* males, respectively, compared with *w^1118^* controls (*p*<0.0001 and *p*<0.0001; one-way ANOVA followed by Tukey’s HSD). Whole-body triglyceride levels were further increased in *AkhR^1^; bmm^1^* males compared with *AkhR^1^* males and *bmm^1^* males (*p*<0.0001 and *p*<0.0001 respectively; one-way ANOVA followed by Tukey’s HSD). *n=*7-8 biological replicates. (B) Whole-body triglyceride levels were significantly higher in *AkhR^1^* females compared with *w^1118^* controls; however whole-body triglyceride levels were not significantly different between *bmm^1^* females and *w^1118^* control females (*p*=0.002 and *p*=0.4256 respectively; one-way ANOVA followed by Tukey’s HSD). Whole-body triglyceride levels were further increased in *AkhR^1^; bmm^1^* females compared to *AkhR^1^* females and *bmm^1^* females (*p*=0.0024 and *p*<0.0001 respectively; one-way ANOVA followed by Tukey’s HSD). *n=*8 biological replicates. Black circles indicate the presence of a mutant allele and open circles indicate the lack of a mutant allele. ** indicates *p*<0.01, **** indicates *p*<0.0001, ns indicates not significant; error bars represent SEM.

**Figure 4 - figure supplement 3.**
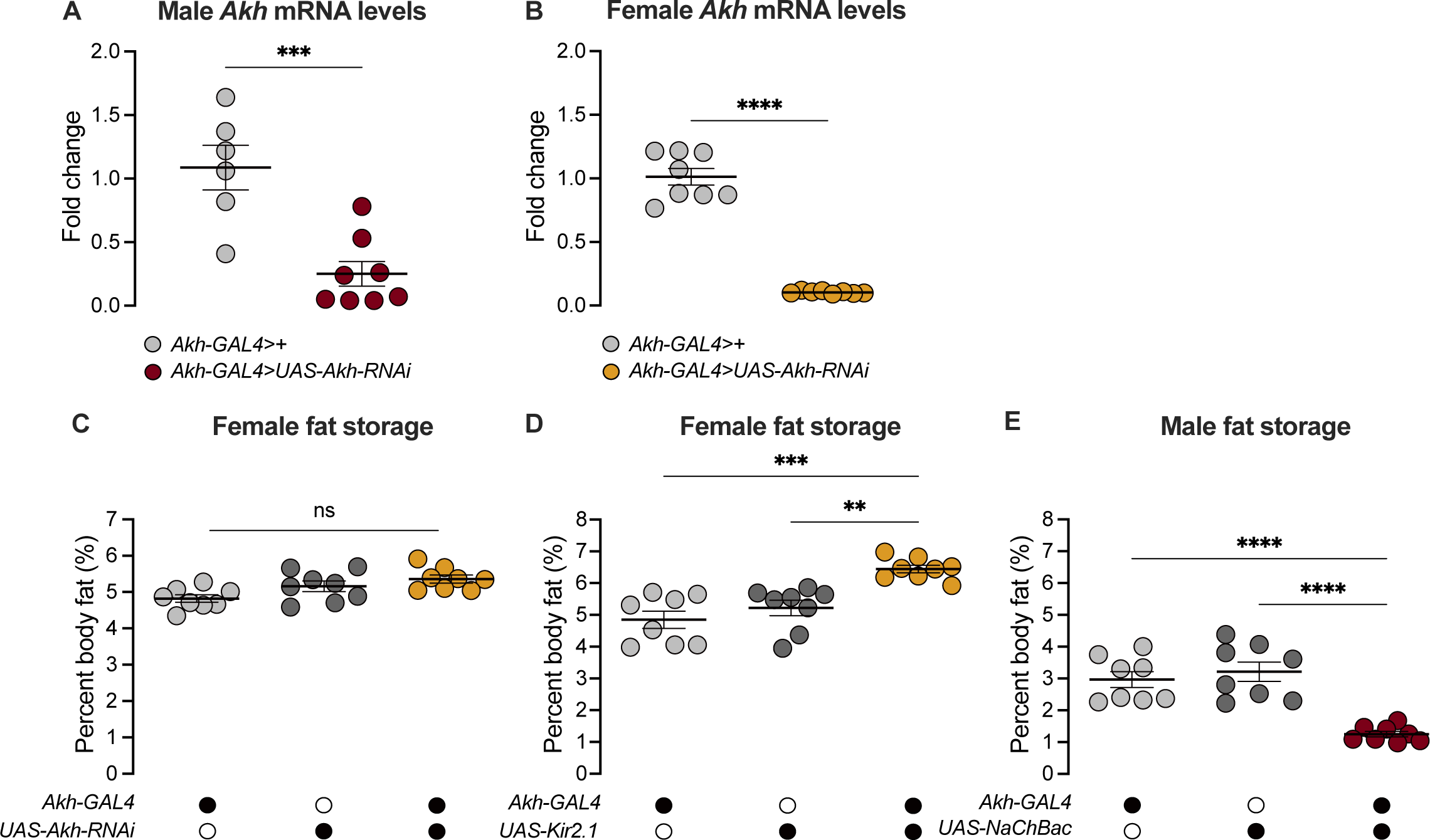
RNAi-mediated Akh knockdown effectively reduced Akh transcripts in both sexes. (A) *Akh* mRNA levels in the head and anterior half of the thorax were significantly lower in *Akh-GAL4>UAS-Akh-RNAi* males compared with *Akh-GAL4>+* controls (*p*=0.0008; Student’s *t*-test). *n*=5-8 biological replicates. (B) *Akh* mRNA levels in the head and anterior half of the thorax were significantly lower in *Akh-GAL4>UAS-Akh-RNAi* females compared with *Akh-GAL4>+* controls (*p*<0.0001; Student’s *t*-test). *n*=8 biological replicates. (C) Whole-body triglyceride levels were not significantly different between *Akh-GAL4>UAS-Akh-RNAi* females and both *Akh- GAL4>+* and *+>UAS-Akh-RNAi* controls (*p*=0.0136 and *p*=0.4845 respectively; one-way ANOVA followed by Tukey’s HSD). *n*=8 biological replicates. (D) Whole-body triglyceride levels were significantly higher in *Akh-GAL4>UAS-Kir2.1* females compared with *Akh-GAL4>+* and *+>UAS-Kir2.1* controls (*p*=0.0001 and *p*=0.0022 respectively; one-way ANOVA followed by Tukey’s HSD). *n*=8 biological replicates. (E) Whole-body triglyceride levels were significantly lower in *Akh-GAL4>UAS-NaChBac* males compared with *Akh-GAL4>+* and *+>UAS-NaChBac* controls (*p*<0.0001 and *p*<0.0001 respectively; one-way ANOVA followed by Tukey’s HSD). *n*=8 biological replicates. Due to independent experiments with a shared GAL4 control, *Akh-GAL4>+* males are shared between Figure 4E and Figure 4 – figure supplement 3E. *Akh-GAL4>+* females are shared between Figure 4F and Figure 4 – figure supplement 3D. Black circles indicate the presence of a transgene and open circles indicate the lack of a transgene. ** indicates *p*<0.01, *** indicates *p*<0.001, **** indicates *p*<0.0001, ns indicates not significant; error bars represent SEM.

**Figure 4 - figure supplement 4.**
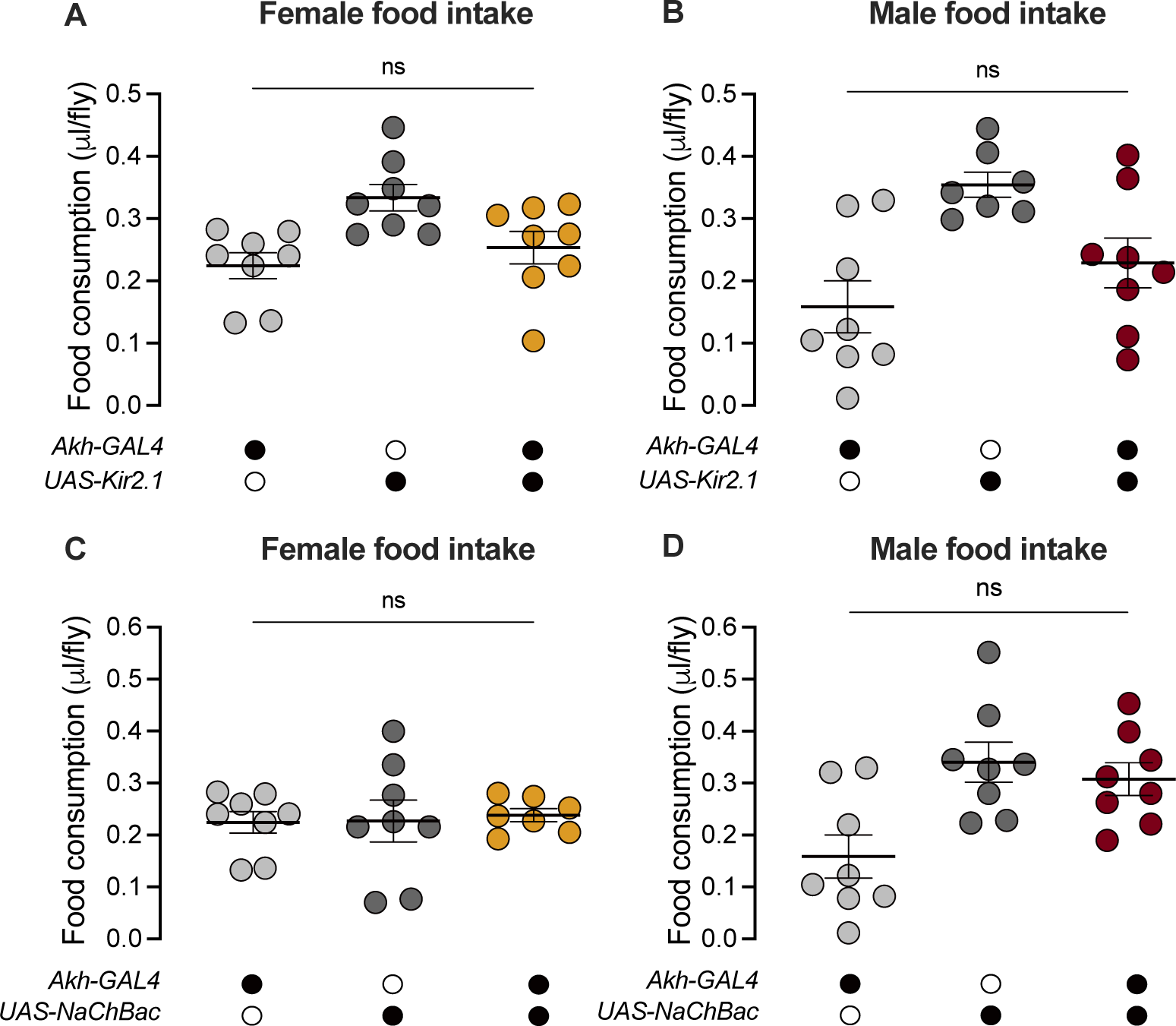
Activity of the Akh-producing cells does not regulate food consumption in either sex. (A) Food consumption was not significantly different between *Akh-GAL4>UAS-Kir2.1* females and *Akh-GAL4>+* and *+>UAS-Kir2.1* controls (*p*=0.6488 and *p*=0.0539 respectively; one-way ANOVA followed by Tukey’s HSD). *n*=8 biological replicates. (B) Food consumption was not significantly different between *Akh-GAL4>UAS-Kir2.1* males and *Akh-GAL4>+* and *+>UAS-Kir2.1* controls (*p*=0.3623 and *p*=0.0638 respectively; one-way ANOVA followed by Tukey’s HSD). *n*=7- 8 biological replicates. (C) Food consumption was not significantly different between *Akh-GAL4>UAS-NaChBac* females and *Akh-GAL4>+* and *+>UAS-NaChBac* controls (*p*=0.9369 and *p*=0.9571 respectively; one-way ANOVA followed by Tukey’s HSD). *n*=7- 8 biological replicates. (D) Food consumption was not significantly different between *Akh-GAL4>UAS- NaChBac* males and both *Akh-GAL4>+* and *+>UAS-NaChBac* controls (*p*=0.0266 and *p*=0.8141 respectively; one-way ANOVA followed by Tukey’s HSD). *n*=8 biological replicates. Due to independent experiments with a shared GAL4 control, *Akh-GAL4>+* females are shared between Figure 4 – figure supplement 4A and Figure 4 – figure supplement 4C. *Akh-GAL4>+* males are shared between Figure 4 – figure supplement 4B and Figure 4 – figure supplement 4D. Black circles indicate the presence of a transgene and open circles indicate the lack of a transgene. ns indicates not significant; error bars represent SEM.

**Figure 5 - figure supplement 1.**
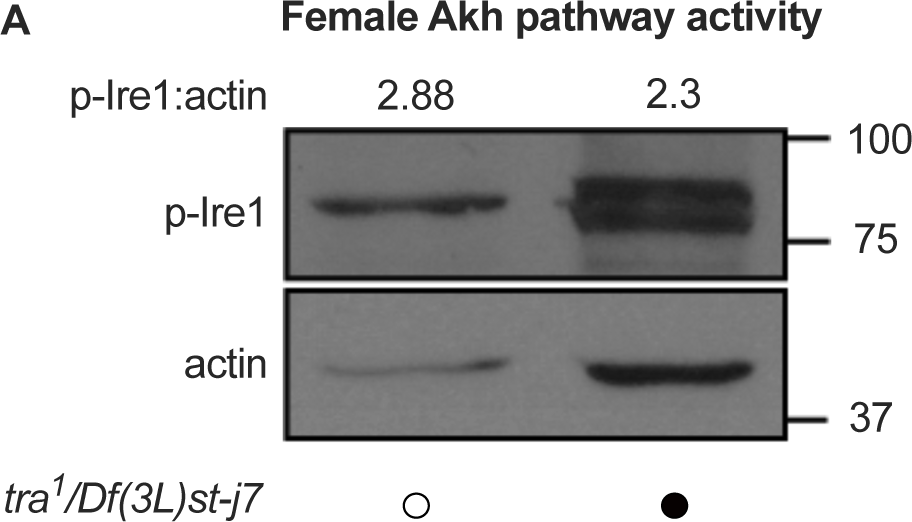
Whole-body p-Ire1 levels in *transformer* mutant flies. (A) Whole-body p-Ire1 levels were not higher in *tra^1^/Df(3L)st-j7* females compared with *w^1118^* control females in one biological replicate. Black circles indicate the presence of a mutant allele and open circles indicate the lack of a mutant allele. Original image for Figure 5 – figure supplement 1A is found in Figure 5 – figure supplement 1 – Source Data 1.

## SOURCE DATA LEGENDS

**Figure 3 – Source Data 1.** Images used to quantify number of Akh-producing cells.

**Figure 3 – Source Data 2.** Images used to quantify neuronal activity of Akh-producing cells.

**Figure 3 – Source Data 3.** Original blots for p-Ire1 and actin in males and females.

**Figure 3 – figure supplement 1 – Source Data 1.** Original blots for p-Ire1 and actin in male vs. female.

**Figure 5 – Source Data 1.** Original blots for p-Ire1 and actin in females with whole-body loss of *transformer*.

**Figure 5- figure supplement 1 – Source Data 1.** Original blots for p-Ire1 and actin in females with whole body loss of *transformer*.

