## Supplementary figures and images for "Sex determination gene *transformer* regulates the male-female difference in *Drosophila* fat storage via the adipokinetic hormone pathway"

### Figure 3 - figure supplement 1B-rep2_actin-pierceplus-6min.png

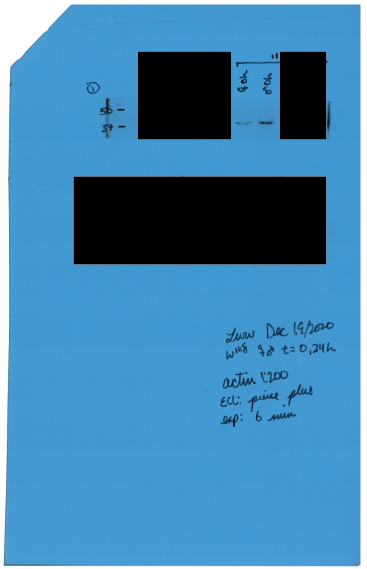

### Figure 3 - figure supplement 1B-rep2_pIRE1-pierceplus-90sec.png

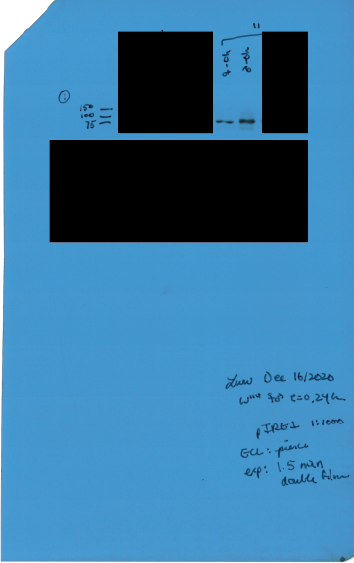

### Figure 3I-rep1_3J-rep4_3K-rep3_actin-pierceplus-6min.png

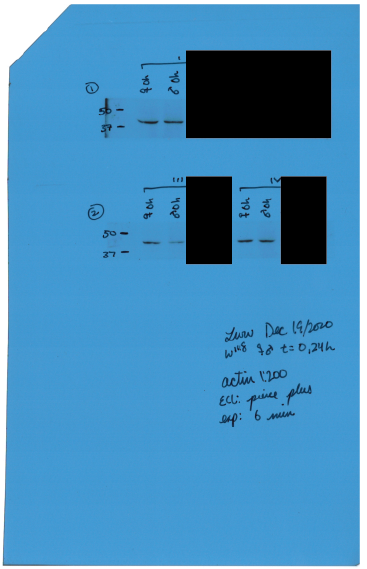

### Figure 3I-rep1_3J-rep4_3K-rep3_pIRE1-pierceplus-90sec.png

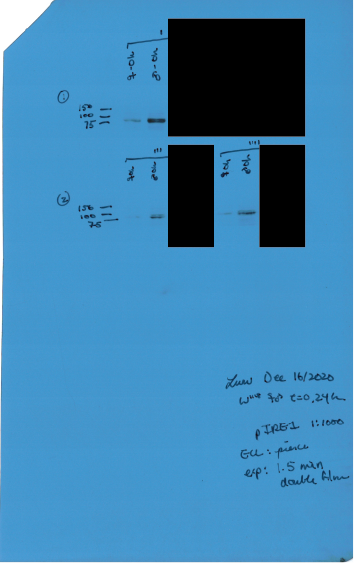

### Figure 5 - figure supplement 1A-rep2_actin-pierce-5min.png

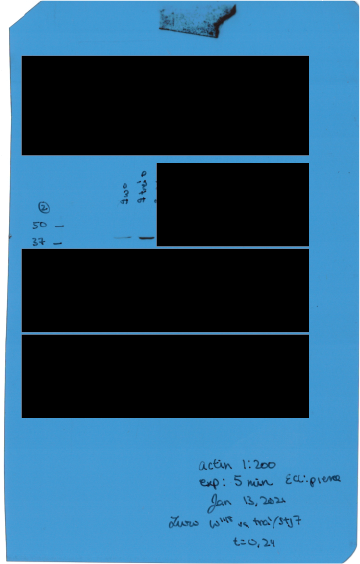

### Figure 5 - figure supplement 1A-rep2_pIRE1-pierce-30sec.png

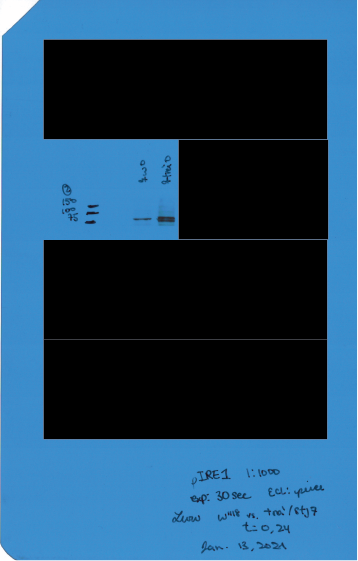

### Figure 5A-rep4_5B-rep3_5C-rep1_actin-pierce-5min.png

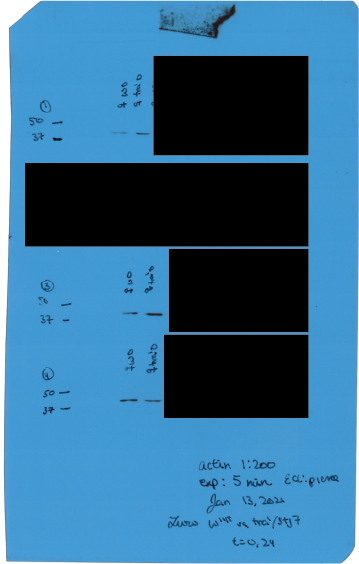

### Figure 5A-rep4_5B-rep3_pIRE1-pierce-1 min.png

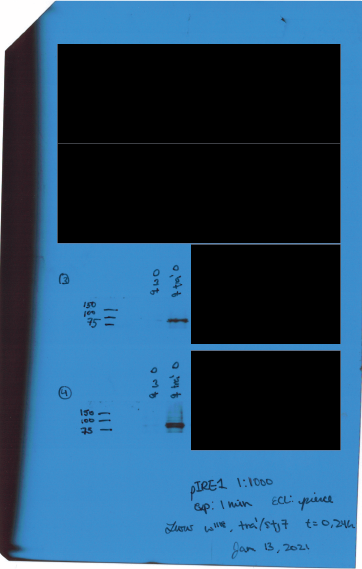

### Figure 5C-rep1_pIRE1-pierce-30sec.png

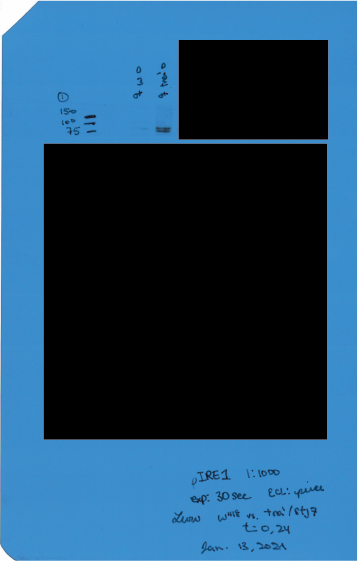
